# SARS-CoV-2 Omicron Envelope T9I adaptation confers resistance to autophagy

**DOI:** 10.1101/2024.04.23.590789

**Authors:** Susanne Klute, Rayhane Nchioua, Arne Cordsmeier, Jyoti Vishwakarma, Lennart Koepke, Hala Alshammary, Christoph Jung, Maximilian Hirschenberger, Helene Hoenigsperger, Jana-Romana Fischer, Fabian Zech, Steffen Stenger, Ruth Serra-Moreno, Ana S. Gonzalez-Reiche, Emilia Mia Sordillo, Harm van Bakel, Viviana Simon, Frank Kirchhoff, Timo Jacob, Dorota Kmiec, Andreas Pichlmair, Armin Ensser, Konstantin MJ Sparrer

## Abstract

To date, five variants of concern (VOCs) of SARS-CoV-2 have emerged that show increased fitness and/or immune evasion. While the continuously evolving escape from humoral immune responses has been analyzed in detail, adaptation of SARS-CoV-2 to human innate immune defenses such as autophagy is less understood. Here, we demonstrate that mutation T9I in the structural envelope (E) protein confers autophagy resistance of Omicron VOCs (BA.1, BA.5 and XBB.1.5) compared to 2020 SARS-CoV-2 or the Delta VOC. Mechanistic analyses revealed that Omicron-associated E T9I shows increased inhibition of autophagic flux and colocalization/interaction with autophagosomes, thus shielding incoming SARS-CoV-2 S pseudotyped virions from autophagy. Rare Omicron isolates carrying ancestral E T9 remain sensitive towards autophagy whereas recombinant early 2020 SARS-CoV-2 expressing E T9I shows increases resistance against autophagy. Our data indicate that the E T9I mutation drives autophagy resistance of the Omicron variants and thus may have contributed to their effective spread.

## INTRODUCTION

The severe acute respiratory syndrome coronavirus 2 (SARS-CoV-2) is the causative agent of the COVID-19 pandemic^1,2^. After crossing the species barrier from bats to humans most likely via a yet unidentified intermediate host at the end of 2019^1^, SARS-CoV-2 continues to adapt as it circulates within its new human host^3^. Thus, several variants of SARS-CoV-2 with increased transmission efficiency and immune evasion, termed variants of concern (VOC) emerged within the past three years, including Alpha, Beta, Gamma, Delta and since late 2021 Omicron^3^. Currently (April 2024), various subvariants of Omicron dominate the pandemic including XBB,EG.5.1, HK.3 and most recently JN.1^4–8^. All Omicron variants are characterized by a large number of mutations in the genome (∼50-100), compared to earlier SARS-CoV-2 strains^9^. These mainly cluster in the gene encoding the surface protein Spike (S)^9^. Most amino acid changes in Omicron S contribute to evasion from humoral immune responses, such as neutralizing antibodies induced by vaccines or previous SARS-CoV-2 infections. This has allowed the Omicron variant to infect and replicate in hosts with pre-existing immunity against prior SARS-CoV-2 variants^10–12^. Thus, escaping from adaptive immunity by altering the surface glycoprotein is a major driver of manifestation of mutations in the genome of successful SARS-CoV-2 variants^3,4^. However, an analysis of 6.4 million SARS-CoV-2 genomes identified four non-Spike mutations within the top 10 mutations in SARS-CoV-2 that are associated with increased viral fitness (ORF1a P3395H, ORF1a K856R, E T9I and ORF9b P10S)^13^. The impact of these mutations is currently not fully understood.

In addition to the adaptive immune system, activation of innate immune defense mechanisms, such as autophagy, was reported to restrict SARS-CoV-2^14–18^. Autophagy is an evolutionary conserved catabolic pathway that has emerged as an integral part of the innate immune system^19^. During selective autophagy, cytoplasmic cargo, including viruses or viral components, is recognized by autophagy receptors such as p62/SQSTM1 and engulfed in LC3B-II-positive double-membrane vesicles called autophagosomes^20,21^. Subsequently, autophagosomes fuse with lysosomes and the cargo along with the receptor is degraded^25^. As such, autophagic flux, i.e. the turnover of autophagosomes, targets incoming virions, viral proteins or budding viral particles for lysosomal degradation^22–26^. To avoid lysosomal degradation, SARS-CoV-2 perturbs autophagy. Infection with SARS-CoV-2 leads to accumulation of p62/SQSTM1 and LC3B-II *in vitro* and *in vivo*, indicating impaired autophagic turnover^15,16^. In depth analyses revealed that at least five of the ∼30 proteins encoded by SARS-CoV-2, including the non-structural proteins Nsp15 and Nsp6, as well as ORF3a, ORF7a and the structural protein Envelope (E), antagonize autophagy^16,27,28^. Whereas Nsp15 reduces autophagic flux, Nsp6 exploits autophagy to promote degradation of immune sensors^16,29^. Expression of the accessory proteins ORF3a and ORF7a resulted in the accumulation of non-acidified autophagosomes in the cytoplasm indicating that they prevent autophagosome turnover^16^. Mechanistically, ORF3a decreases fusion of lysosomes with autophagosomes by interfering with the assembly of the fusion machinery^16,30–32^, whereas ORF7a reduces the acidity of autophago-/lysosomes^16,33^. It was suggested that ORF3a also mediates exploitation of the autophagic machinery to promote SARS-CoV-2 budding^31^. Interestingly, the E protein, one of the four structural proteins of SARS-CoV-2, was shown to inhibit autophagic flux^16^. The 75 amino acid long E protein was reported to assemble into a viroporin consisting of five membrane spanning E proteins and enabling ion channel activity^34,35^. Intracellularly, E was reported to localize to endosomes and lysosomes, trigger ER stress responses and eventually affect host cell survival^36,37^. However, the impact and cellular targets of E in autophagy antagonism are currently unknown.

Here, we show that the mutation T9I in E that emerged with the Omicron lineage of SARS-CoV-2 confers increased resistance against autophagy and protects the incoming virion from autophagy. Mechanistic analyses revealed that T9I increases E localization to autophagosomes as well as interaction with autophagosome-associated SNX12, ST12, TMEM87b and ABCG2. Furthermore, our data shows that entry of viral particles carrying E T9I is less restricted by autophagy. Rare Omicron patient isolates lacking T9I are sensitive towards pharmaceutical autophagy induction, whereas recombinant early 2020 SARS-CoV-2 carrying E T9I gains autophagy resistance. Thus, our results identify escape from autophagy as an evolutionary trajectory of SARS-CoV-2 and reveal E T9I as the mutation allowing Omicron to escape autophagy.

## RESULTS

### The Omicron variant of SARS-CoV-2 is resistant to autophagy

The emergence of the Omicron variant marked a significant shift in SARS-CoV-2 evolution. Earlier strains and VOCs were rapidly outcompeted in late 2021 by successive subvariants of Omicron, such as BA.1, BA.2, BA.5 and XBB, that dominate until today (Fig. 1a). We aimed to understand whether SARS-CoV-2 Omicron adapted to escape not only adaptive immunity, but also innate immunity, in particular autophagy. Therefore, we analyzed the impact of Torin-1, which targets the mTOR complex and activates autophagy^34,35^, on the replication of various VOCs in Calu-3 cells. Our data shows that Torin-1 treatment reduced replication of an early 2020 SARS-CoV-2 isolate (NL-02-2020, here after NL) as well as the Delta variant in a dose-dependent manner by up to 50-fold (Fig. 1b, Extended Data Fig. 1a). In contrast, all tested Omicron variants were largely (BA.1, BA.5) or even fully (XBB.1.5.) resistant to autophagy induction (Fig. 1b). It was previously suggested that inhibition of mTOR promotes SARS-CoV-2 by modulation of the expression of interferon-induced transmembrane proteins (IFITMs)^38^. However, endogenous IFITM levels were unaffected by Torin-1 in Calu-3 cells despite robust autophagy induction as indicated by decreased endogenous SQSTM1/p62 levels and LC3B-I to II conversion (Extended Data Fig. 1b). As revealed by area under the curve analyses, accumulated viral RNA production by BA.1, BA.5 or XBB.1.5. was 8.2-, 5.2- or 13.6-fold less affected by Torin-1 compared to NL (Fig. 1c). Autophagy resistance of Omicron variants was confirmed by determining viral titers 48 h post infection (Fig. 1d, Extended Data Fig. 1c). While infectious viral yields of NL were decreased by almost 4000-fold upon 250 nM Torin-1 treatment, the Omicron subvariants BA.1, BA.5 and XBB.1.5. were restricted by autophagy induction only 20-, 300- or 11-fold (Fig. 1d).

**Figure 1.**
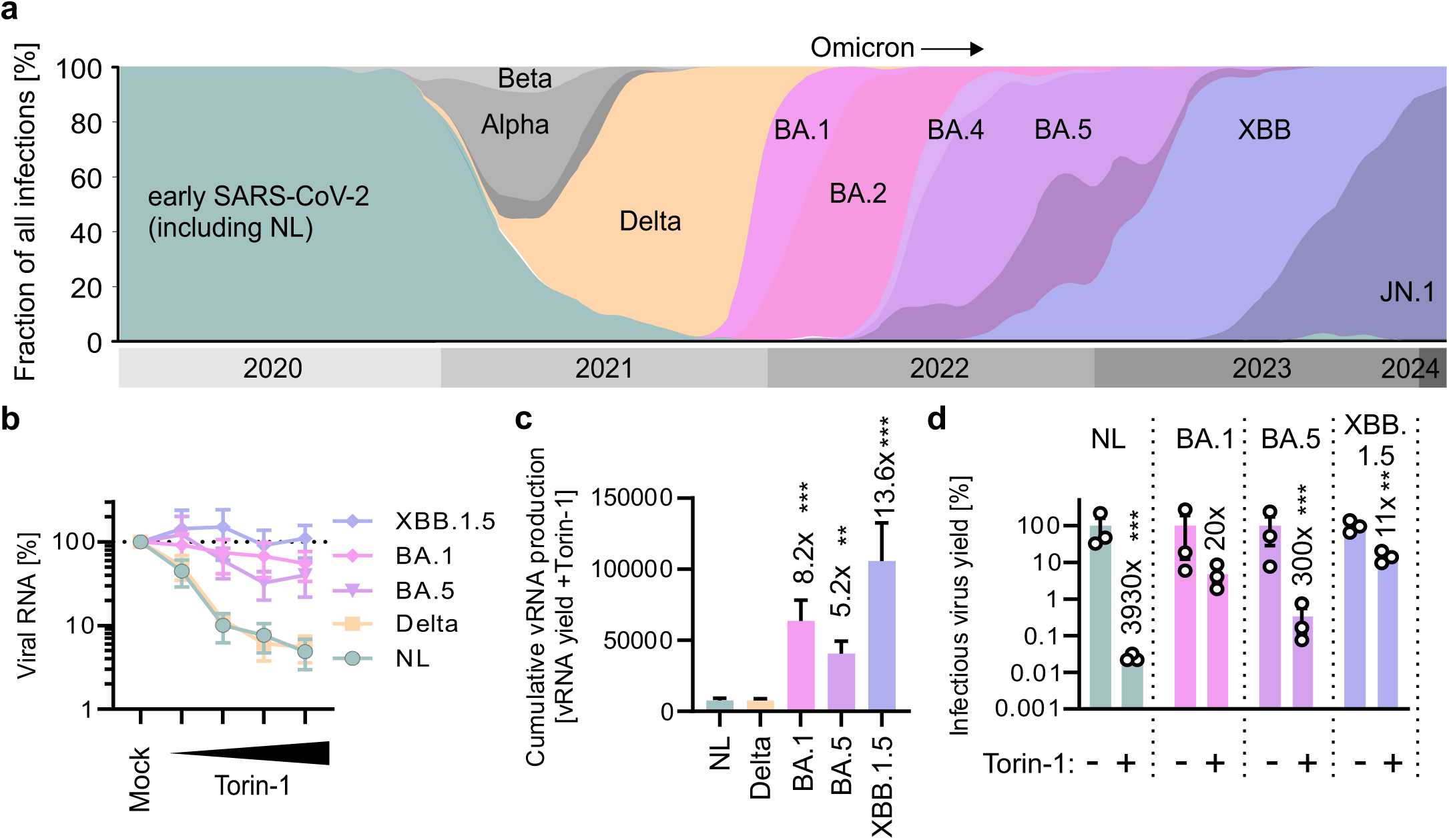
The Omicron VOC is less sensitive against autophagy. **a,** Relative abundance of indicated SARS-CoV-2 strains until 31st December 2023, Data from Nextstrain, retrieved February 2024. **b**, Quantification of SARS-CoV-2 N viral RNA in the supernatant of Calu-3 cells infected with indicated SARS-CoV-2 strains (MOI 0.05) and treated with increasing amounts of Torin-1 (0.016-1 µM) by qPCR 48 h post infection. N = 3-6+SEM. **c**, Area under the curve analysis of the data in (b). N = 3-6+SEM. **d**, Infectious SARS-CoV-2 in the supernatant of Calu-3 cells treated with 250 nM Torin-1 or left untreated and infected with indicated viruses (MOI 0.05) as assessed by TCID50. N = 3±SEM. Student’s t-test with Welch’s correction. **, p<0.01; ***, p<0.001.

These results indicate a pronounced decrease in autophagy sensitivity among Omicron variants compared to ancestral SARS-CoV-2 strains and previous VOCs, indicating acquisition of increased resistance to host cell intrinsic defenses.

### The Omicron-specific T9I mutation in E leads to increased autophagosome accumulation

To investigate the molecular mechanism underlying autophagy evasion by Omicron, we focused on three proteins: ORF3a, ORF7a and E, that were identified by previous studies as key factors encoded by SARS-CoV-2 that antagonize autophagic flux^13,14^. The Delta and the Omicron VOCs harbor distinct consensus mutations (Covariants) in these three genes compared to NL/HU-1: Delta (ORF3a S26L, ORF7a V82A, T120I), BA.1 (E T9I), BA.5 (ORF3a T223I; E T9I), XBB (ORF3a T223I; E T9I, T11A) (Fig. 2a). To determine which of the VOC-specific mutations may alter the sensitivity to autophagy, we introduced single point mutations in NL-derived ORF3a, ORF7a and E by site-directed mutagenesis (Fig. 2b). To analyze the impact of the mutated proteins on autophagy, we employed a flow cytometry-based system^39,40^. In brief, HEK293T cells stably expressing the autophagy marker protein GFP-LC3B stably are mild permeabilized by saponin. Membrane-bound GFP-LC3B, which decorates autophagosomes, remains associated to the cells. Thus, the remaining GFP fluorescence can serve as a proxy for autophagosome content of a cell. Expression of VOC associated SARS-CoV-2 constructs in our autophagy reporter HEK293T cells revealed that all NL-derived and VOC-associated mutant forms of ORF3a and ORF7a caused accumulation of similar levels of autophagosomes (Fig. 2b). However, expression of Omicron-derived E T9I resulted in significantly increased autophagosome accumulation compared to NL-derived E (E WT) or Omicron T11A E (Fig. 2b, right panel). To discern whether autophagosomes levels are increased in the presence of E due to increased autophagic flux or impaired autophagosome turnover, we monitored endogenous LC3B-I to LC3B-II conversion as well as SQSTM1/p62 levels, two hallmarks of autophagy^40^. In the presence of SARS-CoV-2 E LC3B-II and p62 accumulate, suggesting that E inhibits autophagic flux (Fig. 2c). In line with the flow cytometry assay, accumulation of endogenous LC3B-II was increased in the presence of E T9I compared to E WT. Dose-dependency assays revealed that E T9I outperforms accumulation of autophagosomes by E T9 over a broad range of the expression levels (Fig. 2d). To corroborate that E inhibits autophagosome turnover, we impaired autophagosome-lysosome fusion using saturating concentrations of Bafilomycin A1^40^. The impact of both E variants on cellular autophagosome levels as assed by GFP-LC3B flow cytometry is decreased in the presence of Bafilomycin A1, further confirming that E inhibits flux^40^ (Fig. 2e). Finally, accumulation of autophagosomes in the presence of E T9I compared to E T9 was confirmed by quantifying GFP-LC3B-positive puncta (=autophagosomes) in HeLa cells transiently expressing WT E or E T9I (Fig. 2f).

**Figure 2.**
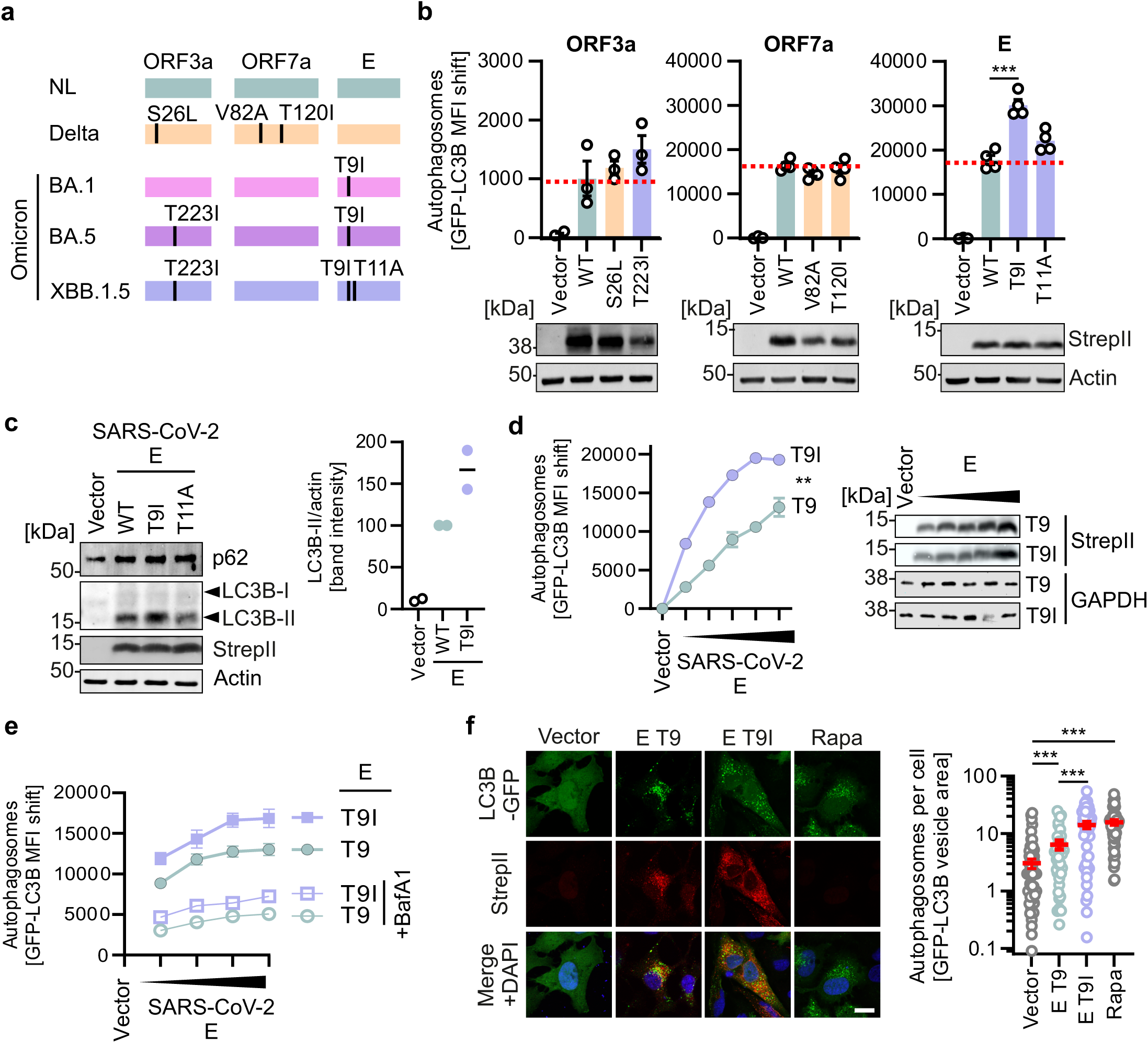
Mutation T9I enhances autophagy antagonism of E. **a**, Schematic depiction of the major autophagy antagonists of SARS-CoV-2, ORF3a, ORF7a and E (Envelope). Specific VOC-associated mutations compared to early 2020 SARS-CoV-2 are annotated. **b**, Quantification of autophagosome levels by flow cytometry in HEK293T autophagy reporter cells (HEK293T-GL) transiently expressing StrepII-tagged SARS-CoV-2 proteins (48 h post transfection). N = 3±SEM. (Top panels). Student’s t-test with Welch’s correction. ***, p<0.001. Immunoblots stained with anti-StrepII and anti-GAPDH. (Bottom panels) **c**, Immunoblots of HEK293T-GL cells overexpressing Omicron-specific E mutants were stained with anti-p62, anti-LC3B, anti-StrepII and anti-Actin (left panel). Quantification of LC3B-II/Actin band intensities of the immunoblots detecting Omicron-specific E mutants (right panel). N=2. **d**, Quantification of autophagosome levels by flow cytometry in HEK293T autophagy reporter cells (HEK293T-GL) transiently expressing increasing amounts of StrepII-tagged SARS-CoV-2 E (48 h post transfection). N = 3±SEM. (Left panel). Exemplary immunoblots stained with anti-StrepII and anti-GAPDH. (Right panels) **e**, Quantification of autophagosome levels by flow cytometry in HEK293T-GL cells transiently expressing increasing amounts of StrepII-tagged SARS-CoV-2 E (48 h post transfection) and treated with 250 nM Bafilomycin A1 (empty dots) or medium (filled dots) 4 h before harvest. N = 4±SEM. **f**, Exemplary confocal immunofluorescence images of HeLa-GFP-LC3B (green) cells transiently expressing StrepII tagged SARS-CoV-2 E T9 and E T9I (red) or treated with Rapamycin (Rapa; 1 µM, 24 h) (scale bar, 10 µm) (left panel). Quantification of the number of autophagosomes (=GFP-LC3B positive puncta) in the images in the left panel. N=35-110±SEM. (right panel). Student’s t-test with Welch’s correction. **, p<0.01; ***, p<0.001.

In summary, these results indicate that the mutation T9I increases the ability of SARS-CoV-2 E to antagonize autophagic flux. Notably, E T9I is the only mutation common to all Omicron subvariants but absent in previous SARS-CoV-2.

### Impact of T9I mutation on E pore and intracellular localization

The E protein of SARS-CoV-2 assembles as a pentameric alpha-helical complex in lipid bilayers, forming an ion channel permissive for Ca^2+^ that can also transport K^+^, Na^+^ in a pH-dependent manner^41^. While T9 is not directly part of the pore-forming core transmembrane helix it is located at the beginning (mouth) of the transmembrane helix (Fig. 3a)^35^. To examine the impact of T9I on pore opening, we performed molecular dynamics modeling leveraging a previously published NMR structure of E (PDB: 7K3G)^40^. This approach allowed us to stimulate the dynamic behavior of the protein upon introducing T9I and evaluate any changes in pore size or conformation. However, this analysis suggests that the T9I mutation has a minimal impact on pore size, with the wild-type E T9 exhibiting a pore size of 14.17Å, whereas the mutant E T9I shows only a slight reduction to 13.61Å (Figs. 3b, c, Extended Data Fig. 2a). To assess whether the viroporin function of E affects its role in autophagy, we employed a E viroporin inhibitor (BIT225)^43^. Treatment with non-cytotoxic concentrations of BIT225 had no impact on autophagosome accumulation levels in the presence of E in HEK293T GFP-LC3B autophagy reporter cells (Extended Data Fig. 2b, c). Previous data showed that E mainly localizes to intracellular vesicles such as lysosomes, late endosomes and autophagosomes^16,37^. Confocal microscopy of HeLa-GFP-LC3B cells transiently expressing either the T9 or T9I E, co-stained with antibodies against endogenous LAMP1, Rab7a and LC3B, showed that both variants co-localize with LAMP1-positive lysosomes, Rab7a-positive late endosomes, and LC3B-positive autophagosomes (Fig. 3d-f). However, while E T9I showed a similar localization to lysosomes as E T9 (Fig. 3d), it was significantly less present on late endosomes, but increasingly co-localized with autophagosomes (Figs. 3e, f).

**Figure 3.**
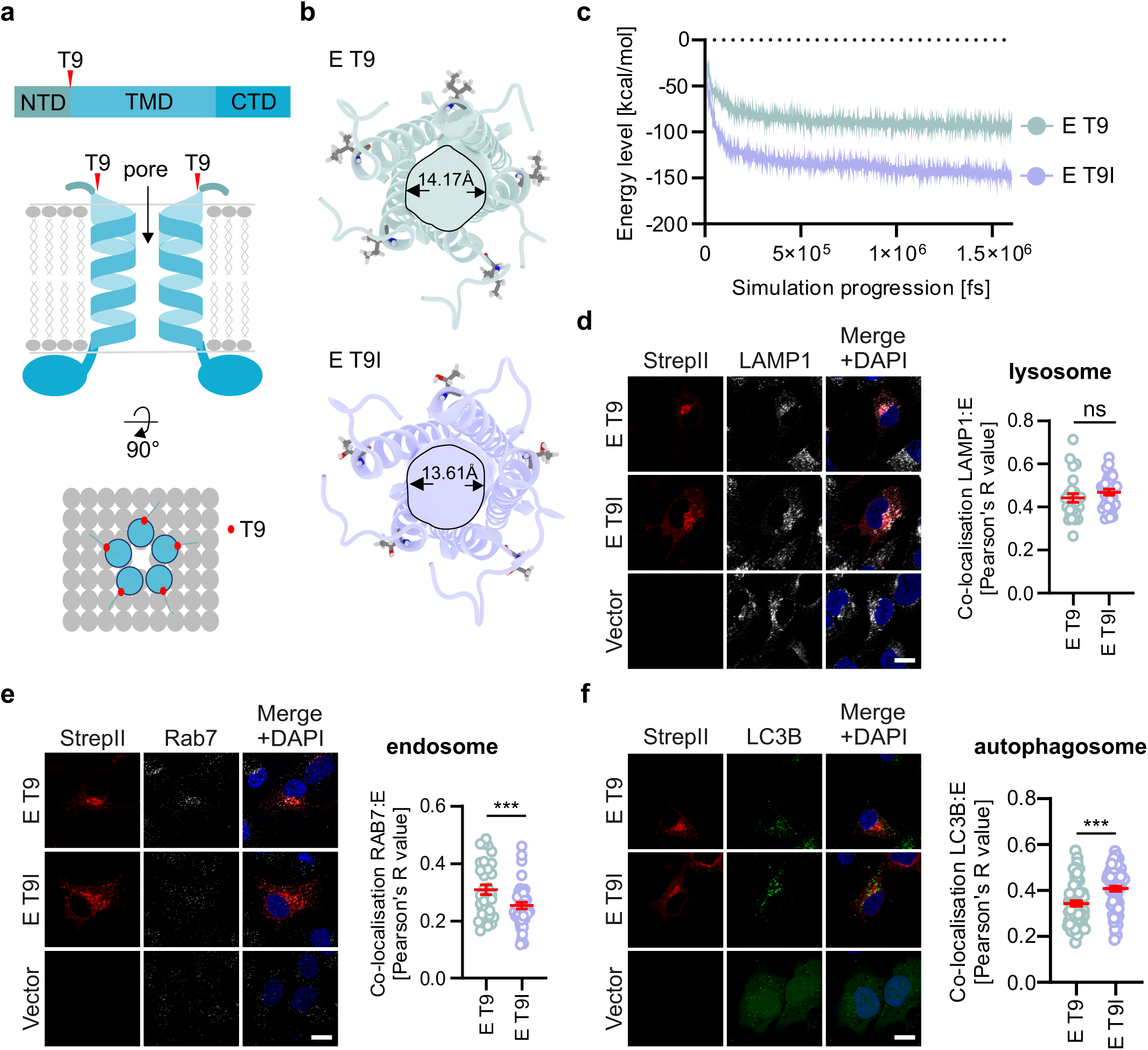
Impact of mutation T9I on E viroporin assembly and intracellular localization. **a,** Schematic depiction of the structure of SARS-CoV-2 E. NTD, N-terminal domain. TMD, Trans-membrane domain. CTD, C-terminal domain. The position of T9 in E is highlighted in red. **b**, Schematic representation of the E protein ion channel depicted as a cartoon showing a transmembrane view. The ninth amino acid is highlighted as a stick, with the colors representing carbon (grey), oxygen (red) and hydrogen (white). The diameter of E protein ion channels is depicted in Ångström (Å). **c,** Exemplary energy curve of the reactive molecular dynamics simulation for SARS-CoV-2 E T9 and E T9I. **d-f**, Exemplary confocal scanning microscopy images of HeLa-GL cells transiently expressing StrepII-tagged SARS-CoV-2 E (red), co-stained with anti-LAMP1 (white) (d), anti-Rab7 (white) (e), anti-LC3B (green) (f). Scale Bar, 10 µm. DAPI, nuclei (blue). Quantification of the co-localization between E variants and indicated co-stained markers using Pearson’s correlation. N=26-73±SEM. Student’s t-test with Welch’s correction. ***, p<0.001. ns (non-significant).

Taken together, these data suggest that the T9I mutation in E does not affect its viroporin assembly, but may shift its localization towards autophagosomes.

### T9I increases E interactions with autophagy-associated proteins

To investigate the impact of T9I of the cellular interactome of E, we performed a differential interactome analysis via large-scale pulldowns of E. To this end, we constructed A549 cells stably expressing either T9 or T9I E and purified the proteins along with their cellular interaction partners. Subsequently, the co-purifying proteins were identified by mass spectrometry (Supplementary Table 1). Principal component analysis revealed that the interactomes of E T9 and E T9I cluster differentially indicating altered primary interaction partners (Fig. 4a). Next, we used Gene ontology (GO) analysis to determine the biological processes associated with the proteins interacting with E T9I. Our results indicate that proteins enriched in E T9I (fold change >2 and p<0.005) are associated with biological processes like Endosomal Transport (GO:0016197) Endocytic Recycling (GO:0032456), Vesicle-Mediated Transport To The Plasma Membrane (GO:0098876) Retrograde Transport, Endosome To Golgi (GO:0042147) (Extended Data Fig. 3a, Supplementary Data 2). Vulcan plot analyses of the aggregated replicates showed that five proteins (STX12, SNX12, TMEM87b, ABCG2 and TAB1) were among the most significantly enriched candidates in the E T9I fraction (Fig. 4b, Extended Data Fig. 3a, b). The SNARE protein STX12 regulates protein transport between late endosomes and the trans-Golgi network but was also reported to be required for autophagosome formation^44^. SNX12 regulates cargo sorting in endosomes, but also locates to sites enriched in phosphatidylinositol 3-phosphate that plays a key role in autophagosome assembly^45^. TMEM87b was predicted to be involved in endosome to Golgi retrograde transport^46^. ABCG2 belongs to the superfamily of ATP-binding cassette (ABC) transporters, and was shown to enhance autophagy^47^. Members of the family of TAB proteins, such as TAB2 and TAB3, were reported to inhibit autophagy, however, not TAB1^48^. To determine whether the mutation T9I increases the spatial interaction of E with the five interaction partners we performed proximity ligation assays (PLA) in HeLa cells (Figs. 4c-g, Extended Data Fig. 3c). These data revealed that mutation T9I in E increased localization/recruitment to SNX12, STX12, TMEM87B, and ABCG2, whereas proximity to TAB1 was not affected. To understand whether these interactions are required for the function of E in autophagy, we depleted HEK293T autophagy reporter cells of STX12, SNX12, TMEM87b, ABCG2, or TAB1 using siRNAs. Knockdown efficiency was >90% (Extended Data Fig. 3d). Analysis of the autophagosome levels using flow cytometry showed that depletion of SNX12, STX12, TMEM87b and ABCG2, but not TAB1 nearly fully abrogated autophagosome accumulation induced by expression of E (Fig. 4h). Of note, in most cases E T9I showed slightly enhanced autophagy antagonistic activity even upon depletion of the respective interaction partner indicating more efficient utilization of the cellular binding partners.

**Figure 4.**
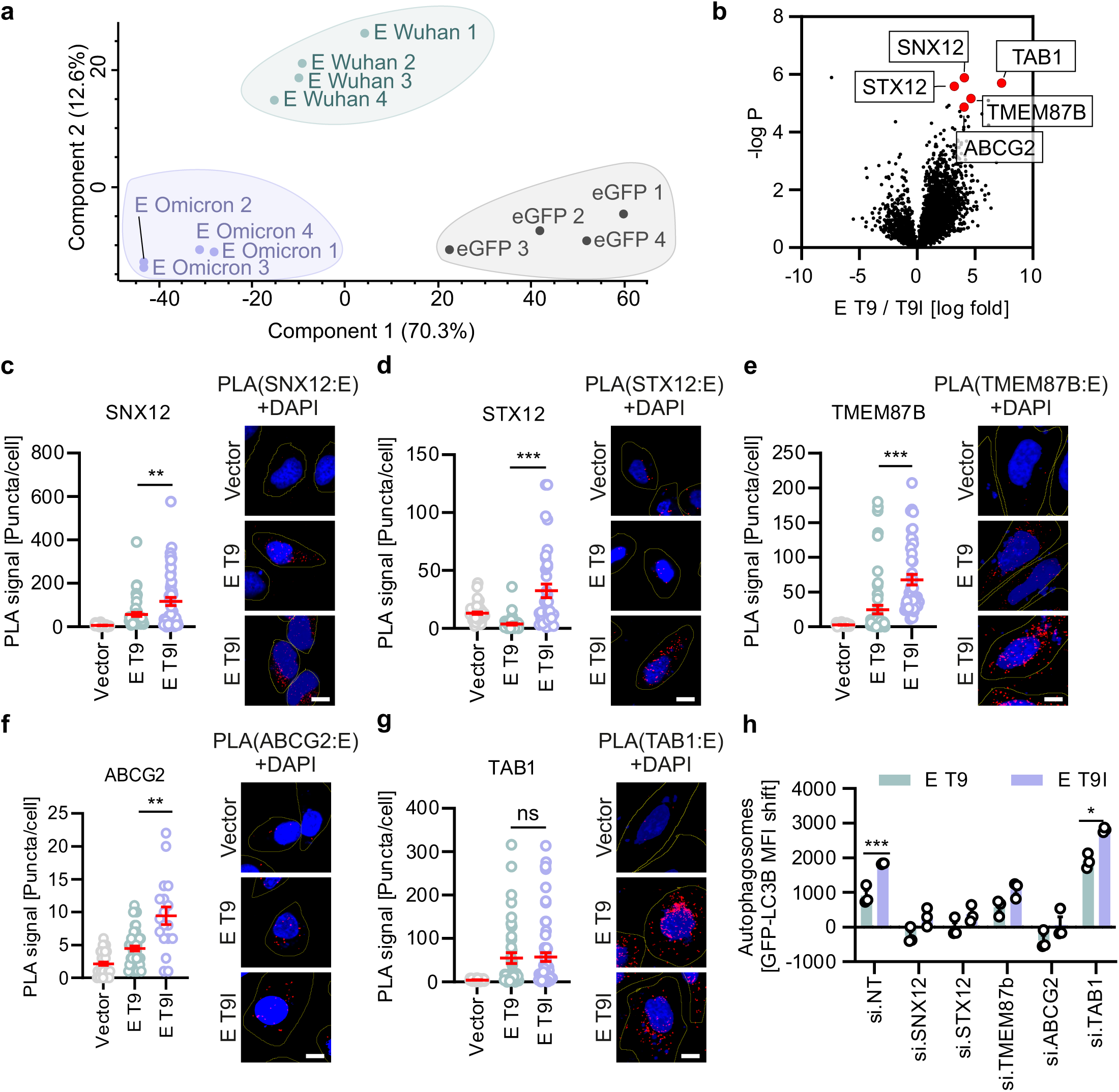
E T9I has increased affinity to autophagosome-associated proteins. **a,** Principal component analysis of the differential interactome data (Supplementary Table 1), the individual replicates are separated (black: GFP controls, Green: E T9 pulldown, Purple: E T9I pulldown) **b,** Volcano plot of the differential interactome analysis showing enriched proteins in E T9I pulldown versus the P value (-log P). Five highly significantly enriched proteins are highlighted in red and via labels**. c-g,** Quantification of proximity ligation assays between transiently expressed SARS-CoV-2 E variants 48 h post transfection in HeLa cells and endogenous SNX12, STX12, TMEM87b, ABCG2 and TAB1, as indicated. Exemplary images depicted. PLA signal, red. Scale Bar, 10µm. DAPI, nuclei (blue). N=18-59±SEM. **h**, Quantification of autophagosome levels by flow cytometry in HEK293T autophagy reporter cells (HEK293T-GL) transiently expressing StrepII-tagged SARS-CoV-2 E variants (48 h post transfection) and depleted of indicated proteins by siRNA. N = 3±SEM. Student’s t-test with Welch’s correction. *, p<0.05; **, p<0.01; ***, p<0.001.

Taken together, these results suggest that E T9I interacts more strongly with components of autophagosome assembly machinery (SNX12, STX12, TMEM87b and ABCG2) and that the presence of these interaction partners is required for E-mediated autophagosome accumulation.

### E T9I does not impact virion composition or production but increases autophagy resistance

Autophagy may target incoming virions for lysosomal degradation and thus restrict viral entry^22,25^. Thus, only structural proteins, unlike non-structural proteins, may promote autophagy evasion upon entry. To examine the impact of E on virion entry separately from replication, we used a Vesicular Stomatitis Virus (VSV)-based pseudoparticles (VSVpp), that is well-established to recapitulate major steps of SARS-CoV-2 entry^12,49,50^. To this end, we pseudotyped VSV that expressed GFP instead of G with SARS-CoV-2 S and either T9 E or T9I E (Extended Data Fig. 4a). Expression of SARS-CoV-2 E and S in producer HEK293T cells infected with VSVΔG resulted in the production of virions that contain the VSV proteins and both SARS-CoV-2 E and S (Fig. 5a). E T9 and E I9 were incorporated with similar efficiency into VSV particles (Extended Data Fig. 4c). The presence or absence of E had no significant effect on SARS-CoV-2 S and VSV M levels in the supernatants indicating similar efficiency of particle production in the presence or absence of autophagy (ATG5 KO cells) (Figs. 5a, Extended Data Fig. 4b-d). Similar S incorporation levels into VSV were confirmed by SARS-CoV-2 S ELISA (Fig. 5b). Previous studies suggested that E may affect functionality and processing of virion-associated S^34^. However, western blot analyses showed that S in the supernatant was processed similarly in the presence of both E T9 or T9I (Fig. 5a). This was confirmed by assessing the ACE2-S interaction *in vitro* using ELISA. S-containing VSV pseudoparticles produced in the presence of E T9 and T9I interacted with recombinant ACE2 at a similar efficiency (Fig. 5c). To assess the impact of autophagy on incoming virions, we treated Caco-2 cells with Torin-1 before the infection. As expected, autophagy induction reduced infection with single-round VSVpp-Spike by about 50% (Fig. 5d, Extended Data Fig. 4e). Inclusion of E T9 into the particle only marginally altered autophagy resistance of the pseudoparticles. In contrast, VSV pseudoparticles carrying E T9I were almost completely resistant towards Torin-1 treatment (Fig. 5d). In line with this, infection with S-pseudotyped particles carrying E T9I was most efficient in WT MRC5 cells compared to particles carrying E T9 particles. However, the advantage of having E T9I in the virion was lost in autophagy-negative MRC5 cells (ATG5KO) (Fig. 5e).

**Figure 5.**
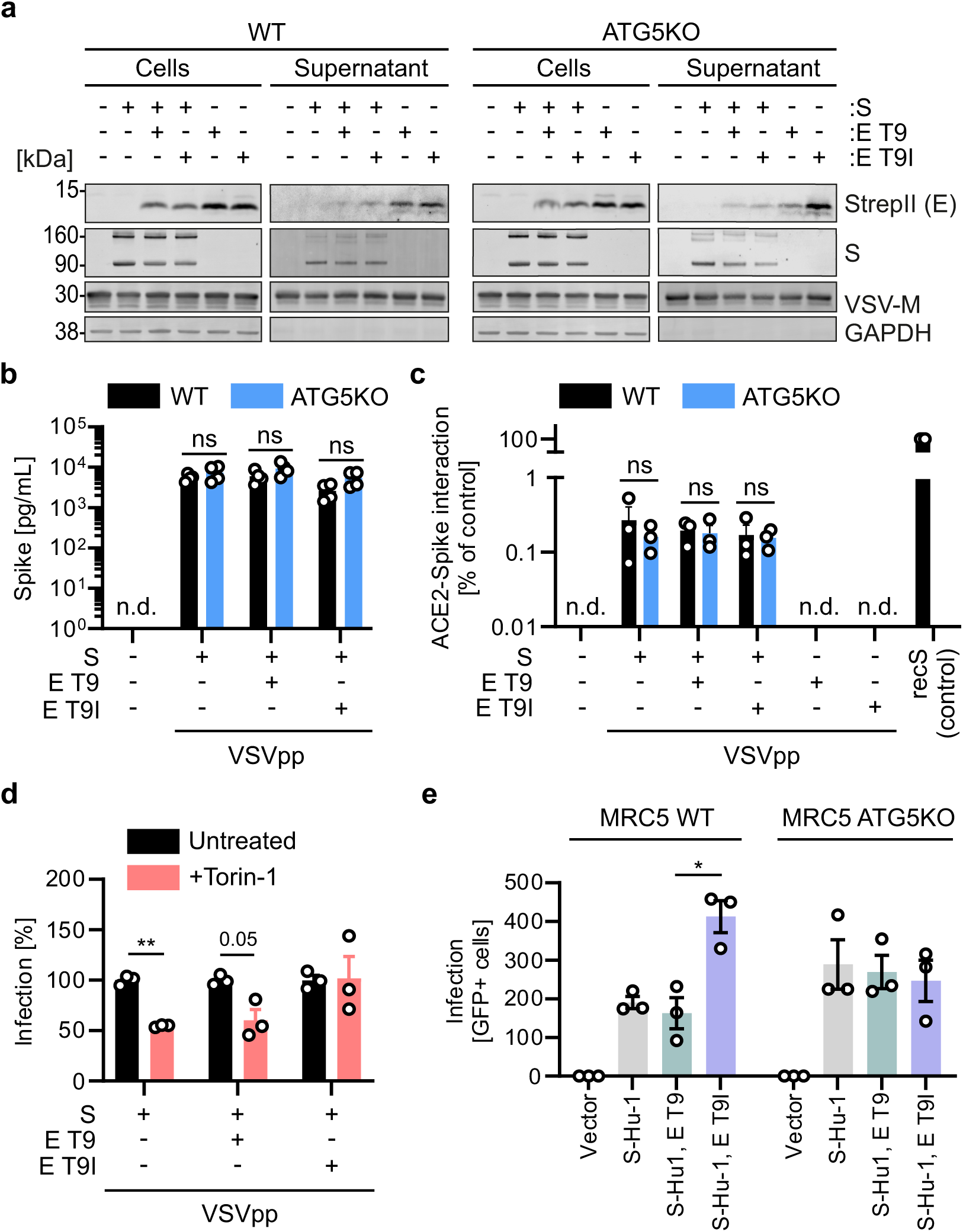
VSV pseudoparticles carrying E T9I are less sensitive towards autophagy. **a,** Exemplary immunoblots of whole cell lysates and supernatants of HEK293T WT (left panel) and ATG5 KO HEK293T (right panel) cells transiently expressing SARS-CoV-2 S, E T9 or E T9I as indicated and infected with VSV-ΔG-GFP (MOI 3). Blots were stained with anti-StrepII, anti-V5 (Spike), anti-VSV-M and anti-GAPDH. **b,** Concentration of S in VSVpp containing supernatants of HEK293T WT and ATG5 KO cells in (a) assessed by Spike-ELISA. N=4±SEM. **c,** ACE2 binding of VSVpp produced on HEK293T WT and ATG5 KO cells as in (a) as assessed by a Spike-ACE2 interaction ELISA. recS, recombinant Spike. N=3±SEM. **d,** Analysis of the infectivity of the particles produced in (a) by dividing the number of infected Caco-2 cells, either mock or Torin-1 (4 h, 0.5 µM) treated (S4e) with the amount of Spike (b). Infection was normalized to 100% for mock. **e**, Infection of MRC5-ACE2 WT and ATG5 KO cells with VSV(GFP)ΔG pseudoparticles produced in (a) containing the indicated proteins. Infected GFP+ cells were automatically quantified after 22 h post infection. N = 3±SEM. Student’s t-test with Welch’s correction. *, p<0.05; **, p<0.01. ns (non-significant). n.d. (non detectable).

Collectively, these results suggest that the incorporation of E T9I in VSV pseudotyped with SARS-CoV-2 S does not alter particle assembly or intrinsic infectivity but promotes autophagy resistance.

### Rare naturally-occurring Omicron variants with E T9 are autophagy sensitive

Coinciding with the emergence of the Omicron VOC in late November 2021, E T9I rapidly became predominant in the circulating SARS-CoV-2 strains (Figs. 6a, b). Of note, E T9I was only sporadically (<1%) present in previous variants circulating in the pre-Omicron era (Fig. 6b, data from Nextstrain, January 2024). In 2023, 97.28% of the sequences available of circulating SARS-CoV-2 encoded the E I9 substitution, while only 0.76% retained the T at position 9, with 1.95% of the sequences showing undefined amino acid at position 9 (Fig. 6c, data from CovSpectrum, January 2024^51^). The Mount Sinai Health System (MSHS) is one of the largest health care providers in the New York City metropolitan area. This health system supports a well-integrated pathogen surveillance infrastructure enabling real time monitoring of pathogens detected in people receiving care at one of the eight hospitals as well as at any of the many outpatient clinics. We utilized the ongoing surveillance of circulating SARS-CoV-2^52–54^ to validate the data from Nextstrain, findings from the Mount Sinai Pathogen Surveillance (PSP-MS) database, which contains over 10,000 SARS-CoV-2 genomes generated between March 2020 and December 2023, revealed that prior to the emergence of the Omicron variant, the predominant variant across all lineages was the wild-type E T9. Only rare SARS-CoV-2 genomes encoded the E T9I mutation, observed in early B.1.* lineages (Supplementary Data 3). However, with the advent of the Omicron variant, a notable shift occurred: the majority of Omicron lineages encode E T9I mutation (Fig. 6b, Extended Data Fig. 5a). Only eight SARS-CoV-2 Omicron isolates retain the ancestral E9T (Supplementary Data 4). To explore the phenotype of this substitution in the context of replication competent authentic SARS-CoV-2 viruses, we cultured two closely related isolates Omicron BA.2 with E T9I (USA/NY-MSHSPSP-PV58179/2023, EPI_ISL_12711111) (Fig. 6d, Extended Data Fig. 5a) and with E T9 (USA/NY-MSHSPSP-PV58079/2023, EPI_ISL_12711042) (Fig. 6d, Extended Data Fig. 5a). Comparison of the replication of these two viruses on Calu-3 cells, infected at the same MOI, showed that PV58179 (E T9) grew to approximately 10-fold higher titers than the PV58079 (E T9I) isolate (Fig. 6e). Consistent with the pseudotyping experiments, treatment with Torin-1 during the infection reduced the replication of T9 E PV58179 more than 4-fold whereas autophagy activation did not impact T9I E PV58079.

**Figure 6.**
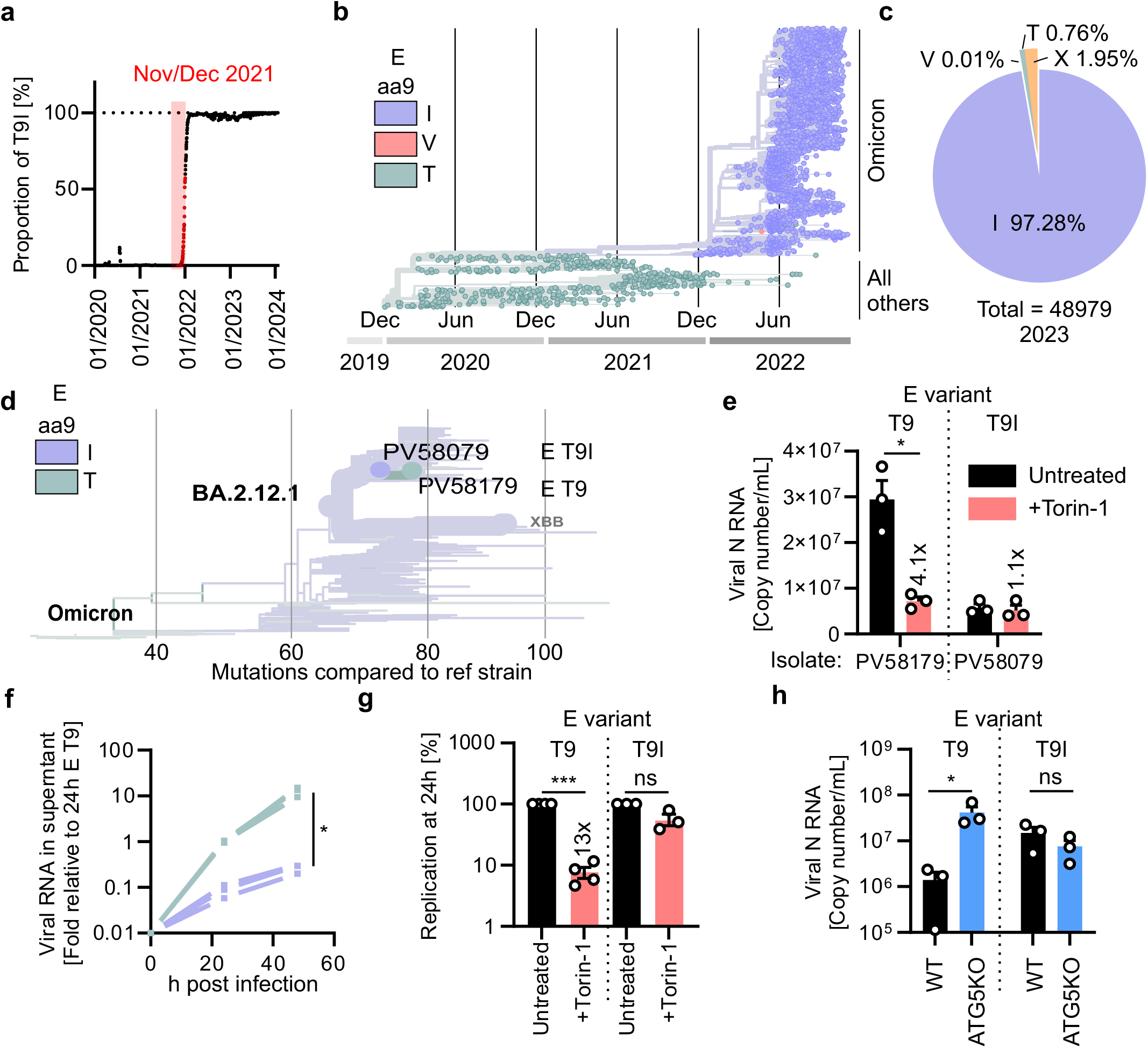
T9I E promotes autophagy resistance of patient-isolated and recombinant SARS-CoV-2. **a,** Ratio of E T9I mutation in all SARS-CoV-2 strains sampled from 01/2020 to 07/2023. Nov/Dec 2021 highlighted in red. **b,** Phylogenetic tree of SARS-CoV-2 variants since early 2020, sourced from Nextstrain, November 2023. Amino acid at position 9 in E is highlighted by colors. **c,** Percentages of amino acid residues at position 9 in E of 1365370 Omicron strains. **d**, Excerpt of a phylogenetic tree showing the collection of SARS-CoV-2 isolates from the Mount Sinai surveillance program. The relation between E T9 coding PV58079 and E T9I coding PV58179 isolates is highlighted. **e**, Absolute quantification of viral RNA in the supernatant of Calu-3 cells mock treated or treated with Torin-1 (250nM) and infected with SARS-CoV-2 patient isolates PV58079 or PV58179 (MOI 0.05) as assessed by qPCR 48 h post infection. N=3±SEM. **f**, Relative quantification of viral RNA in the supernatant of Calu-3 cells infected with rSARS2-E T9 or rSARS2-E T9I as assessed by qPCR 24h and 48h post infection. 24h rSARS2-E T9 is set to 1. N=4±SEM. **g**, Relative quantification of viral RNA in the supernatant of Calu-3 cells treated or non-treated with Torin-1 (250nM) and infected with rSARS2-E T9 or rSARS2-E T9I (MOI 0.05) as assessed by qPCR 24h post infection. N=4±SEM. 24 h mock treated values are set to 100%. N=3±SEM. **h**, Quantification of viral RNA in MRC5 WT or ATG5KO cells by qPCR at 72 h post infection with SARS2-E T9 or rSARS2-E T9I (MOI 0.05). N=3±SEM. Student’s t-test with Welch’s correction. *, p<0.05; ***, p<0.001. ns (non-significant).

These data with authentic viruses suggest that the mutation T9I in E limits replication but augments resistance to autophagy.

### T9I in E increases 2020 SARS-CoV-2 resistance against autophagy

To confirm that E T9I confers resistance against autophagy, we generated recombinant SARS-CoV-2 harboring E T9I in an ancestral 2020 SARS-CoV-2 background (Extended Data Fig. 5b). Both viruses (rSARS2-E-T9 and rSARS2-E-T9I) were rescued on Caco-2 cells^55^. Growth analyses revealed that rSARS2-E-I9 replicated to ∼10-50-fold lower titers than rSARS2-E-T9 (Fig. 6f). rSARS2-E-T9 was highly sensitive towards autophagy induction by Torin-1, reducing replication after 24 h by more than 10-fold (Fig. 6g). In contrast, despite replicating much slower rSARS2-E-I9 was almost completely resistant against autophagy (Fig. 6g). Compared to autophagy incompetent MRC5 cells (ATG5KO MRC5), replication of rSARS2-E-T9 was reduced significantly in WT MRC5 cells (Fig. 6h). rSARS2-E-T9I replicated to comparable levels in both WT and ATG5KO MRC5 cells as indicated by similar intracellular viral RNA expression (Fig. 6h). This suggests that lack of autophagy does not convey an advantage for rSARS2-E-T9I.

In summary, the E T9I mutation confers resistance against autophagy to recombinant SARS-CoV-2 at the price of reduced replication capacity.

## DISCUSSION

Within the last three years SARS-CoV-2 adapted to its new human host giving rise to variants characterized by increased infectivity and immune escape^1,3,9,56,57^. Here, we show that a single point mutation in the E protein (T9I) of the Omicron VOCs conveys autophagy resistance.

While our analyses show that E T9I promotes autophagy resistance of the virions, it may come at a cost - reduced replication fitness *in vitro* (Fig. 6). This translates to ∼10-fold lower replication rates of recombinant SARS-CoV-2 carrying E T9I in a Hu-1 background but it is also apparent by increased replication fitness of rare patient-derived isolates of Omicron BA.2 carrying the original T9 residue in E. This could be a contributing factor to the delayed emergence of the E T9I mutation, despite its sporadic presence in pre-Omicron SARS-CoV-2 isolates^53,54^. It is tempting to speculate that acquiring E T9I contributed to the slower replication of early Omicron strains^18,56,58^. Of note, more recent Omicron subvariants show increased replication competence; thus, it is likely that they evolved compensatory mutants that promote replication^6,58,59^. The identity of these compensatory mutations is, however, unknown.

Why would a VOC with mutations reducing replication speed *in vitro* emerge? Superior immune escape, not replication speed, was suggested to be a defining factor in the success of early Omicron subvariants^11,18,60^. A previous study analyzing 6.4 million SARS-CoV-2 genomes shows that E T9I - as one of the few non-Spike mutations - conveyed a fitness advantage in the population^13^. This suggests that on a population level E I9 had an advantage *in vivo* despite its attenuated replication in *in vitro* experiments. It is remarkable that evasion of autophagy may be more important than a 10-fold reduction in replication fitness, and contributes to the evidence that immune escape may be more important for coronaviruses *in vivo* than fast replication.

Mechanistically, E T9I increasingly localizes to autophagosomes (Fig. 3f) and interacts more strongly with autophagosome-associated factors STX12, SNX12, TMEM87b, ABCG2 than E T9 (Fig. 4b). Our data further reveals that these factors are required for the impact of E on autophagy (Fig. 4h). Of note, all four targets are endosome/lysosome associated proteins^46,61^. STX12 and SNX12 are part of the SNARE complex and together with SNAP29 mediate fusion of autophagosomes with lysosomes^44,62^. It thus seems likely that E may impair the function of the fusion machinery, resulting in impaired autophagic flux^63^. However, future studies are required to clarify the precise impact of E on these four proteins. In addition, it needs to be clarified, whether counteraction of autophagy is the only function of E that has been altered by mutation T9I.

Our data suggests that the pore function of E is not required for its function in autophagy and the pore size is not affected by mutation T9I. This was a bit surprising since viroporins frequently affect autophagy due to their impact on the pH of intracellular vesicles or via ER stress^64,65^. E was shown to be capable of transporting K+, Na+, and Ca2+ ions across membranes^37,41,66^. Of note, while not impacting autophagy, the pore of E activates other parts of innate immunity, such as the inflammasome^28,67^. Blockage of the E channel activity reduces the inflammatory response towards SARS-CoV-2 infection, but also reduces viral replication^28,43,66,67^. Thus, it had been suggested that E may prevent the premature activation of S, by alkalizing the ERGIC^37^. However, the precise pro-viral roles of the ion channel function of E remain to be determined.

Our data shows that T9I in E may enhance escape from autophagy upon cell entry. SARS-CoV-2, in addition to fusing at the plasma membrane, may enter cells through the endosomal route^50,68,69^. Thus, incoming virions may be redirected to autophagy for lysosomal degradation^22,24,70^. In addition, E may facility endosomal escape of virions to ensure the genomic information makes it unscathed to the cytoplasm to establish the infection. Interestingly, it was suggested the Omicron variants, depending on the tissue and cell type show increased entry via the endosomal route^71–73^. It is thus tempting to speculate that E T9I promotes evasion of incoming virions by autophagy, enabling increased entry via the endosomal route. Unfortunately, it is currently little understood how incoming virions are targeted by autophagy, and our understanding of how this mechanism contributes to the induction of immune defenses by increasing abortive infections remains limited. The COVID-19 pandemic allowed unprecedented insights into the adaptation of zoonotic viruses to their new human host^28,57^. Thus, one may ask whether E T9I is an evolutionary adaptation to human autophagy? Notably, autophagy is highly evolutionarily conserved across all eukaryotes from yeast to humans^19,22,23,74^. The immune system of the reservoir species of SARS-CoV-2, bats, is well-known to be highly tolerant of viruses without developing significant diseases^75^. Of note, it was shown that, as opposed to other mammals, ageing bats show increased levels of homeostatic autophagy^76^. However, it remains unknown whether autophagy in bats is also uniquely regulated, despite high sequence similarity of core components of the autophagic machinery. Bat Coronaviruses closely related to SARS-CoV-2 all share a T in position 9 in E. Future analysis of autophagy escape by bat coronaviruses may yield important insight on differences between bat and human autophagy and thus may both enhance our understanding of the evolution of autophagy and bats as a unique virus reservoir species.

In addition to the evolving escape from adaptive immunity, adaptation to innate immune defenses such as autophagy, may be a major contributing factor in the current transition of SARS-CoV-2 to endemic status^1,77,78^. Infections with Omicron can overcome adaptive immunity induced by infection with previous variants or early vaccines^59,73^. Adjusted Omicron-specific vaccines as well as surviving an Omicron infection again confer a protective adaptive response. However, unlike adaptive immune responses, our innate defenses are a rigid defense system allowing for little adaptation to changing viruses. Thus, gaining innate immune resistance may represent a more permanent advantageous evolutionary step for SARS-CoV-2.

In summary, our data shows that the Omicron associated T9I mutation in E confers increased resistance towards autophagy, suggesting an evolutionary adaptation to autophagy of SARS-CoV-2. Reduced sensitivity to autophagy in combination with reduced sensitivity to IFN as well as neutralizing antibodies from prior infections and vaccinations may have contributed to the rapid spread of the Omicron variant.

## MATERIALS AND METHODS

### Cell culture and viruses

All cells were cultured at 37 °C under a 5% CO_2_ atmosphere and 95% relative humidity. HEK293T (ATCC, #CRL3216), Vero E6 (ATCC, #CRL-1586), and ATG5 KO HEK293T^62^ were cultivated in Dulbecco’s Modified Eagle Medium (DMEM) (Gibco, #41965039) containing 10% (v/v) heat-inactivated fetal bovine serum (FBS) (Gibco, #A5256701), 10 mg/ml gentamicin (PAN-Biotech, #15710-049), and 2 mM L-glutamine (PAN-Biotech, #P04-80100). The construction of autophagy reporter HEK293T and HeLa cells stably expressing GFP-LC3B (GL) was reported previously^63^. HEK293T-GL and HeLa-GL cells were cultivated in DMEM supplemented with 10% (v/v) heat-inactivated FBS, 10 mg/ml gentamicin, and 2 mM L-glutamine. Caco-2 cells (kindly provided by Prof. Holger Barth, Ulm University, Ulm, Germany) were maintained in DMEM supplemented with 20% (v/v) heat-inactivated FBS, 10 mg/ml gentamicin, and 2 mM L-glutamine. Calu-3 cells (kindly provided by Prof. Manfred Frick, Ulm University, Ulm Germany) were cultivated Minimum Essential Medium Eagle supplemented (MEM, Sigma-Aldrich, #M4655) with 10% (v/v) heat-inactivated FBS (during viral infection) or 20% (v/v) FBS (during all other times), 100 mg/ml streptomycin, 100 U/ml penicillin (PAN-Biotech, #P06-07100) and 2 mM L-glutamine. Vero E6 cells (BEI Resources, #NR-54970) overexpressing ACE2 and TMPRSS2 were maintained in DMEM supplemented with 10% (v/v) heat-inactivated FBS, 1% (v/v) of 100x MEM Non-Essential Amino Acids (NEAA) (Gibco, #11140050), 100 mg/ml streptomycin, 100 U/ml penicillin, 2 mM L-glutamine and 1 mM sodium pyruvate (Thermo Fisher, #11360039), 100 µg/ml of Normocin (InvivoGen, #ant-nr-1) and 3 µg/ml of Puromycin (Invivogen, #ant-pr-1). ACE2 overexpressing MRC5 cells were cultivated in DMEM supplemented with 20% (v/v) heat-inactivated FBS, 10 mg/ml gentamicin, and 2 mM L-glutamine. For ATG5 KO MRC5-ACE2 cells 1 µg/mL Puromycin was additionally added to the medium. Mouse I1-Hybridoma cells (ATCC, #CRL-2700) were cultured in Roswell Park Memorial Institute (RPMI) Medium 1640 (Gibco, #21875-034) with 10% (v/v) heat-inactivated FBS, 100 mg/ml streptomycin, 100 U/ml penicillin, 2 mM L-glutamine. HEK293T cells expressing a T7 polymerase and SARS-CoV-2 N protein^43^ for virus reconstitution were cultivated in DMEM supplemented with 10% (v/v) heat-inactivated FBS, 2 mM Gluta-MAX™ (Thermo Fisher Scientific, #35050061), 25 mM HEPES (Thermo Fisher Scientific, #15630080), 5 µg/mL blasticidin (InvivoGen, #asnt-bl-1), and 2 µg/mL puromycin. HEK293T (ATCC, #CRL-11268) and A549 cell lines for mass spectrometry experiments were cultured in DMEM supplemented with 10% (v/v) heat-inactivated FBS and 1% Penicillin-streptomycin. All cell lines were tested to be mycoplasma free.

The SARS-CoV-2 variants, B.1.617.2 (Delta) and B.1.1.529 (Omicron BA.5), were kindly provided by Prof. Dr. Florian Schmidt and Dr. Bianca Schulte (University of Bonn, Bonn, Germany). The BetaCoV/Netherlands/01/NL/2020 (NL-02-2020) lineage and hCoV-19/Netherlands/NH-EMC-1720/2021, lineage B.1.1.529 (Omicron BA.1), were obtained from the European Virus Archive. SARS-CoV-2 XBB1.5 and BA.2 variants (USA/NY-MSHSPSP-PV58179/2023, EPI_ISL_12711111) and (USA/NY-MSHSPSP-PV58079/2023, EPI_ISL_12711042) were banked, sequenced and cultured by the Mount Sinai Pathogen Surveillance Program (Mount Sinai Hospital, New York, USA).

The VSV(GFP)ΔG*VSV-G stock was kindly provided by Prof. Karl-Klaus Conzelmann (LMU Munich, Munich, Germany).

### Propagation of WT SARS-CoV-2

SARS-CoV-2 was propagated on different cell lines depending on the viral variant: Vero E6 (NL-02-2020, Delta), Calu-3 cells (Omicron BA.1), or Vero E6 cells overexpressing ACE2 and TMPRSS2 (XBB 1.5 and BA.2 variants). To this end, 70-90% confluent cells in 75 cm² cell culture flasks were inoculated with the SARS-CoV-2 isolate (multiplicity of infection (MOI) of 0.03-0.1) in 3.5 ml serum-free medium (MEM, Sigma, #M4655). The cells were incubated for 2 h at 37 °C, before adding 20 ml medium containing 15 mM HEPES (Carl Roth, #6763.1). Virus stocks were harvested as soon as strong cytopathic effect (CPE) became apparent. The virus stocks were centrifuged for 5 min at 1,000 g to remove cellular debris, aliquoted, and stored at -80 °C until further use.

### Replication competent authentic SARS-CoV-2 isolates

Residual nasopharyngeal swab specimens from patients with COVID-19 were banked by the Mount Sinai Pathogen Surveillance Program after the completion of the diagnostic process as reported previously^48,49^. Replication-competent SARS-CoV-2 viruses were cultured by inoculating Vero-E6-TMPRSS2-ACE2 cells with 200 μl of viral transport media from the nasopharyngeal swab specimen. The culture media were supplemented with 2% heat-inactivated FBS, 100 μg/ml normocin (Invivogen, #ANT-NR-2), and 0.5 μg/ml amphotericin B (Gibco, #15290026), and maintained for a maximum of 10 days. Upon the appearance of cytopathic effects, culture supernatants were collected, clarified by centrifugation (3,739g for 5 min) and sequence verified. These experiments were performed in a BSL-3 biosafety laboratory at the Icahn School of Medicine at Mount Sinai.

### Generation and propagation of a rSARS-CoV-2

Recombinant SARS-CoV-2 were generated based on the bacmid^64^. In brief, the area encoding for E was replaced via homologous recombination by a kanamycin resistance cassette with *AscI* restriction enzyme sites at each end (Primers dE-Asc-KanS-FP: Aat taa agt tcc aaa cag aaa aac taa tat aat att tag ttc gtg gcg cgc cat gac gac gat aag tag gg and dE-Asc-KanS-RP: Tag cgt gcc ttt gta agc aca agc tga tga gta cga act tgg cgc gcc aac caa tta acc aat tct gat tag; IDT Ultramers, Coralville, ID, USA). Subsequently, the kanamycin resistance cassette could be removed by restriction digestion and a respective PCR fragment containing the E T9I mutation and with overlapping ends was introduced by Gibson assembly using the NEBuilder HiFi DNA Assembly Master Mix (New England Biolabs, #E2621). This PCR fragment was amplified using cDNA from a patient sample containing the E T9I mutation with the primers Efwd gcacaagctgatgagtacgaactt and Erev gaaaaactaatataatatttagttcg. The resulting bacmid was transformed in E. coli and recovered by DNA isolation. The correct assembly was verified by Next Generation Sequencing of the full bacmid. The recovery of infectious virus was performed by transfection of HEK293T T7/N cells and passaging on Caco-2 cells. The method has been described previously^65^.

### Analyzing growth and autophagy sensitivity of rSARS-CoV-2

Calu-3 cells were seeded 24 h prior to infection (3 x 10^4^ per well of a 96-well plate). The cells were treated with 250 nM Torin-1 (EZSolution, #2353) 2 h before infection. After infection with an MOI of 0.05, 6 h post infection the medium was replaced by medium containing 250 nM fresh Torin-1. Viral supernatant was collected at 0 h, 24 h and 48 h post infection and heat inactivated. The supernatants were digested with proteinase K (Sigma-Aldrich, #3115828001) and the relative viral load was determined by qRT-PCR as described previously^64^.

### Impact of autophagy induction on the replication of SARS-CoV-2 strains

1.5 x 10^5^ Calu-3 cells/well were seeded in 24-well plates. For autophagy induction, 24 h post-seeding cells were either left untreated or were stimulated with increasing amounts of Torin-1 (0.016, 0.063, 0.25 or 1 µM). Two hours post-treatment, Calu-3 cells were infected with the indicated SARS-CoV-2 strains using a MOI of 0.05. Six hours post-infection, the cells were washed once with Dulbecco’s Phosphate Buffered Saline (DPBS) (Gibco, #14190-094) and supplemented with fresh medium only or medium containing increasing amounts of Torin-1 (0.016, 0.063, 0.25, 1 µM) as indicated in the figures. Supernatants were harvested at 6 h (for wash control) and 48 h post-infection for qRT-PCR and Tissue culture infectious dose 50 (TCID50) analysis.

### TCID50 titration

1.5 x 10^4^ Vero E6 cells/well were seeded in 96-well F-bottom plates in 100 µl medium and incubated overnight. SARS-CoV-2 stocks or infectious supernatants were serially diluted and 100 µl of the dilution were added to the cells (final dilutions 1:10^1^ to 1:10^10^). The cells were incubated for at least 5 days and monitored for cytopathic effects (CPE). TCID50/mL was calculated according to the Reed-Muench method.

### qRT-PCR

Total RNA of the supernatants collected from SARS-CoV-2 infected Calu-3 cells 48 h post-infection were isolated using the QIAamp Viral RNA Mini Kit (Qiagen; #52906) according to the manufacturer’s instructions. To determine the SARS-CoV-2 N (nucleocapsid) levels quantitative real-time PCR (RT-qPCR) was performed as previously described^22^ using TaqMan Fast Virus 1-Step Master Mix (Thermo Fisher, #4444436) and an OneStepPlus Real-Time PCR System (96-well format, fast mode). The Primers as well as the N primer probe for RT-qPCR were from Biomers (Ulm, Germany) and had the following sequence: Forward primer (HKU-NF): 5’-TAA TCA GAC AAG GAA CTG ATT A-3’; reverse primer (HKU-NR): 5’-CGA AGG TGT GAC TTC CAT G-3’; primer probe (HKU-NP): 5’-FAM (6-carboxyfluorescein)-GCA AAT TGT GCA ATT TGC GG-TAMRA (6-carboxytetramethylrhodamine)-3’. Synthetic SARS-CoV-2 RNA (Twist Bioscience, #102024) was used as a quantitative standard to determine viral copy numbers. All PCR reactions were prepared in duplicates.

RNA of siRNA transfected cells were isolated using the Quick-RNA Miniprep Kit (Zymo Research, #R1055) according to the manufacturer’s instructions. To determine the KD efficiency, reverse transcription and qRT-PCR was performed in one step using the SuperScript III Platinum Kit (Thermo Fisher Scientific, #11732088) on a StepOnePlus Real-Time PCR System (Applied Biosystems) according to the manufacturer’s instructions. TaqMan probes for each individual KD gene and for GAPDH were acquired as premixed TaqMan Gene Expression Assays (Thermo Fisher Scientific) and added to the reaction. The following TaqMan primer probes were used: Snx12-FAM-MGB (Thermo Fisher Scientific, #Hs04999580_s1), Stx12-FAM-MGB (Thermo Fisher Scientific, #Hs00295291_m1), Tab1-FAM-MGB (Thermo Fisher Scientific, #Hs00196143_m1), Tmem87b-FAM-MGB (Thermo Fisher Scientific, #Hs00262432_m1), Abcg2-FAM-MGB (Thermo Fisher Scientific, #Hs01053790_m1) and GAPDH-VIC-TAMRA (Applied Biosystems, #4310884E). Expression level for each target gene was calculated by normalizing against GAPDH using the ΔΔCT method.

### Impact of autophagy induction on endogenous IFITM levels

0.7 x 10^6^ Calu-3 cells/well were seeded in 6-well plates. For autophagy induction, 24 h post-seeding cells were left untreated or were stimulated with increasing amounts of Torin-1 (0.016, 0.063, 0.25 or 1 µM). As positive control for upregulation of IFITM expression, further cells were treated with 1000 U IFN-β (R&D Systems, #8499-IF-010/CF). 24 h post-treatment whole-cell lysates for SDS-PAGE and immunoblotting were prepared.

### Cloning and origin of expression constructs

Plasmids coding for SARS-CoV-2 E-StrepII (pLVX-EF1alpha-SARS-CoV-2-E-2xStrep-IRES-Puro), SARS-CoV-2 M-StrepII (pLVX-EF1alpha-SARS-CoV-2-M-2xStrep-IRES-Puro), SARS-CoV-2 ORF7a-StrepII (pLVX-EF1alpha-SARS-CoV-2-ORF7a-2xStrep-IRES-Puro) and ORF8-StrepII (pLVX-EF1alpha-SARS-CoV-2-ORF8-2xStrep-IRES-Puro) were a kind gift from Nevan Krogan^66^. A plasmid coding for SARS-CoV-2 E 9I-StrepII was generated using pLVX-EF1alpha-SARS-CoV-2-E-2xStrep-IRES-Puro. To this end, the template vector was linearized with the restriction enzymes EcoRI-HF (NEB, #R3101L) and BamHI-HF (NEB, #R3136L). All site directed mutagenesis was performed using the NEBuilder HiFi DNA Assembly Master Mix according to the manufacturer’s instructions. For T9I in E the primers E-T9I_F (GTG TCG TGA GGA TCT ATT TCC GGT GAA TTC GCC GCC ACC ATG TAC AGC TTC GTA TCA GAA GAA ATT GGG ACA CTG ATCG; Biomers) and E-T9I_R (CGA TCA GTG TCC CAA TTT CTT CTG ATA CGA AGC TGT ACA TGG TGG CGG CGA ATT CAC CGG AAA TAG ATC CTC ACG ACA C; Biomers) were used. SARS-CoV-2 E 11A-StrepII was generated using the template pLVX-EF1alpha-SARS-CoV-2-E-2xStrep-IRES-Puro and the primers E-T11A_F (AGA AAC CGG GGC TCT GAT CGT AA; Biomers) and E-T11A_R (TCT GAT ACG AAG CTG TAC; Biomers). SARS-CoV-2 E 9I 11A-StrepII was generated using pLVX-EF1alpha-SARS-CoV-2-E 9I-2xStrep-IRES-Puro as template and the primers E-T9I+T11A_F (AGA AAT TGG GGC TCT GAT CGT AAA TTC; Biomers) and E-T9I+T11A_R (TCT GAT ACG AAG CTG TAC; Biomers). ORF7a 120I-StrepII or ORF7a 82A-StrepII were generated by using pLVX-EF1alpha-SARS-CoV-2-ORF7a-2xStrep-IRES-Puro as a template and primers ORF7a-T120I_F (AAA AGA AAG Atc GAG CTC GAA GGC G; Biomers) and ORF7a-T120I_R (CAG TGT AAA GCA CAA TGT G; Biomers) or primers ORF7a-V82A_F (GCT CGA TCT GCC TCC CCC AAA CTG TTC ATA C; Biomers) and ORF7a-V82A_R (CCT GAG CTG GTA CAC GTG; Biomers). pLVX-EF1alpha was constructed using Q5 Site-Directed Mutagenesis (NEB, #E0554S) according to the manufacturer’s instructions with pLVX-EF1alpha-SARS-CoV-2-ORF8-2xStrep-IRES-Puro (codon-opt.HUM) as template and pLVX-EF1alpha-empty-F (ACG CGT CTC GAG GGA TCC CGC CCC TCT CCC TC; Biomers) and pLVX-EF1alpha-empty-R (GCT AGC GCG GCC GCG AAT TCA CCG GAA ATA GAT CCT CAC) as primers. ORF3a 26L-StrepII and 223I-StrepII were cloned by Q5 Site-Directed Mutagenesis with pTWIST_SARS-CoV-2-ORF3a-2xStrep as template and the primers ORF3a-S26L_F (TGC TAC ACC TCT CGA CTT CGT CAG; Biomers) and ORF3a-S26L_R (TCC TTG ATC TCG CCC TGC) or primers ORF3a-T223I_F (AGC ACC GAC ATC GGC GTC GAG; Biomers) and ORF3a-T223I_R (CAGCTGTGTGCTATACAGCTGGTAGTAA; Biomers). pTwist-Empty Vector was constructed using Q5 Site-Directed Mutagenesis, pTwist_EF1a_3xFLAG_opt TRIM3^45^ as template and the primers pTwist-empty-F (GCT TCC GCC TCC GCC GCT T; Biomers) and pTwist-empty-R (GCT AGC TTG ACT GAC TGA GAT ACA GCG TAC CTT; Biomers). pCG_SARS-CoV-2-Spike-Wuhan-1 was previously described^11^. pTWIST_SARS-CoV-2-ORF3a-2xStrep (codon-opt.HUM) was purchased from Twist Bioscience.

### Generation of S and E containing VSV-pseudoparticles

To produce pseudotyped VSV(GFP)ΔG particles, HEK293T WT and ATG5 KO cells were transfected with Spike (HU-1), Envelope (Hu-1) or Envelope T9I expression constructs (in total 3 µg DNA/well) in 6-well format by using per well 10 µL 1x polyethyleneimine-hydrochlorid (PEI) (Sigma-Aldrich, #764965-1G, 1 mg/mL in H_2_O) and 500 µL Opti-MEM reduced serum media (Gibco, #31985047). In brief, 250 µL Opti-MEM were mixed with 10 µL 1x PEI and incubated for 5 min. Secondly, 250 µL Opti-MEM were mixed with 3 µg DNA. Subsequently, both mixes were combined and incubated for 20 min at RT, then added to the cells. 24h post-transfection the cells were infected with VSV(GFP)ΔG*VSV-G at MOI of 3. 24 h post infection, cells and supernatants containing pseudotyped VSV(GFP)ΔG particles were harvested. Cell debris were removed by centrifugation for 4 min at 500 x g.

### VSV-pseudoparticle assays

6 x 10^3^ Caco-2 cells or MRC5-ACE2 (WT and ATG5 KO) cells were seeded in 384-well plates in 25 µL medium. On the next day, the cells were treated with 10 µL Torin-1 (final concentration on cells 0.5 µM) or medium for 4 h and afterwards infected with 35 µL of supernatant containing VSV(GFP)ΔG particles with S and E. Residual particles carrying VSV-G were blocked before by adding 10% (v/v) of I1 hybridoma supernatant (I1 mouse hybridoma supernatant from CRL-2700, ATCC) to the supernatant. GFP-positive cells were automatically counted 22 h post-infection by using Cytation 3 microplate reader (BioTek Instruments).

### Spike ELISA

Supernatants containing VSV(GFP)ΔG particles with S and E were analyzed for their Spike content by using a SARS-CoV-2 (2019-nCoV) Spike Detection ELISA Kit (SinoBiological, #KIT40591). Samples were diluted 1:200 in 1x dilution buffer provided by the Kit. The ELISA was performed according to the manufacturer’s recommendations. Absorbance at 450 nm was detected using the Vmax kinetic microplate reader (Molecular Devices LLC) and the software SoftMax Pro 7.0.3.

### Spike-ACE2 interaction assay

Supernatants containing VSV(GFP)ΔG particles with S and E were layered on a cushion of 20% sucrose (Sigma-Aldrich, #S9378-1KG) in PBS and centrifuged for 90 min at 4°C and 20,817 x g. The pellet was lysed in 12 µL transmembrane lysis buffer (50mM HEPES pH 7.4 (Sigma-Aldrich, #H3375), 150 mM NaCl (Merck, #106404), 1% Triton X-100 (Sigma-Aldrich, #T8787), 5 mM ethylenediaminetetraacetic acid (EDTA) (Sigma-Aldrich, #E9884) supplemented with 1:500 protease inhibitor (Sigma-Aldrich, #P2714) and heated up for 10 min at 95 °C. For analyzing the interaction between Spike of the samples and ACE2 a COVID-19 Spike-ACE2 Binding Assay Kit (RayBioTech, #CoV-ACE2S2-1) was used. In brief, 10 µL of the lysates were mixed with 40 µL of 1x assay diluent buffer (RayBioTech, #CoV-ACE2S2-1) and added to the ACE2 coated wells (RayBioTech, #CoV-ACE2S2-1). After 2 h incubation time with shaking at RT, the wells were washed 3 times with 200 µL 1x wash buffer (RayBioTech, #CoV-ACE2S2-1) and subsequently incubated with 100 µL monoclonal mouse anti-V5 antibody (1:1000, Cell Signaling Technology, #80076) for 1 h at RT under shaking. After washing, wells were incubated with 100 µL HRP-conjugated anti-IgG mouse (1:1000, RayBioTech, #CoV-ACE2S2-1) for 1h under shaking. After subsequent washing, the wells were incubated with 50 µL TMB one-step substrate reagent (RayBioTech, #CoV-ACE2S2-1) for 30 min under shaking in the dark. The reaction was stopped by adding 50 µL stop solution (RayBioTech, #CoV-ACE2S2-1) and the absorbance was detected at 450 nm with a baseline correction of 650 nm by using the Vmax kinetic microplate reader (Molecular Devices LLC) and the software SoftMax Pro 7.0.3.

### Autophagosome measurement by flow cytometry

For autophagosome quantification HEK293T cells stably expressing GFP-LC3B (GL) were used. 4.5 x 10^4^ HEK293T-GL cells were transiently reverse transfected with 200 ng expression vector in 96-well F-bottom plates by using 2 µL 1x PEI/ µg DNA and 17 µL Opti-MEM reduced serum media^63^. The next day, the medium was replaced with 100 µL fresh medium. 48 h post-transfection, the supernatant was removed and the cells were detached using Trypsin/ EDTA 0.05%/ 0.02% (PAN-Biotech; #P10-023100). After adding medium, the harvested cells were washed with DPBS. Autophagosome levels were quantified as previously described^63^ in basal state or after stimulation with 0.2-25 µM BIT225 (kindly provided by Biotron Limited) for 24 h or stimulation with 0.25 µM Bafilomycin A1 (Santa Cruz Biotechnology, #sc-201550) for 4 h. In brief, cells were treated with DPBS containing 0.05% Saponin (Sigma-Aldrich, #47036) for 20 min at 4°C for permeabilization. Subsequently, the cells were washed twice with DPBS to wash out the non-membrane bound GFP-LC3B out of the permeabilized cells and fixated with 4% paraformaldehyde (PFA) (Santa Cruz Biotechnology, #sc-281692). The mean fluorescence intensity (MFI) of membrane-bound GFP-LC3B was then detected by flow cytometry (FACS-Canto II, BD Biosciences). The MFI value of the control was used as baseline and subtracted.

### Immunofluorescence

1x 10^5^ HeLa-GL cells/well were grown on coverslips with 500 µL medium in 24-well plates and one day later transfected with a control vector or expression vectors of SARS-CoV-2 E (Hu-1) or E T9I by using Opti-MEM reduced serum media and TransIT-LT1 Transfection Reagent (Mirus, #MIR2306) according to the manufacturer’s instructions. The supernatant was removed 6 h post-transfection and 500 µL fresh medium was added to the cells. As a positive control, cells were treated with 100 nM Bafilomycin A1 or 1 µM Rapamycin (Merck, #553211-500UG) 24 h before harvesting. 48 h post-transfection the cells were washed twice with DPBS, fixed with 4% PFA for 20 min at RT and blocking and permeabilization was performed by using DPBS with 0.5% Triton-X-100 and 5% FBS for 1 h at RT. After washing with DPBS, the cells were stained with the following primary antibodies diluted in DPBS with 1% FBS for 2 h at 4 °C: Monoclonal mouse anti-StrepII-tag (1:200; Novus Biologicals, #NBP2-43735), monoclonal rabbit anti-LAMP1 (1:200; Cell Signaling, #9091S) and monoclonal rabbit anti-Rab7 antibody (1:500; Abcam, #ab137029). Subsequently, the cells were washed three times with DPBS supplemented with 0.1 % Tween 20 (Sigma-Aldrich, #P7949). In the next staining step, fluorescently labeled secondary antibodies Goat anti-Mouse IgG (H+L) AF568 (1:400; Thermo Scientific, #A21245), Goat anti-Rabbit IgG (H+L) AF647 (1:400; Thermo Scientific, #A11004) as well as DAPI (4’,6-Diamidino-2-Phenylindole, Dihydrochloride) (1:1000; Invitrogen, #D1306) were diluted in DPBS with 1% FBS and incubated for 2 h at 4 °C. Afterwards, the cells were washed three times with DPBS supplemented with 0.1 % Tween 20 and additionally once with deionized water. The coverslips were mounted on microscope slides by using mowiol mounting medium (10% (w/v) Mowiol 4-88 (Carl Roth, #0713), 25% (w/v) Glycerol (Sigma-Aldrich, #G5516), 50% (v/v) Tris-Cl at 0.2 M pre-adjusted to pH 8.5 (AppliChem GmbH, #A2264) and 2.5% (w/v) DABCO (Carl Roth, #0718)) and then allowed to dry overnight at 4 °C. Images were captured using a Zeiss LSM 710 confocal laser scanning microscope with the ZEN imaging software or Leica DMi8 confocal microscope with the LAS X imaging software. Co-localization analysis was performed using the Huygens Professional 19.04 software, and the Pearson coefficients were calculated via the “Huygens Colocalization Analyzer”. Autophagosome counts (GFP-LC3B puncta) per cell were determined using Fiji ImageJ^63^. In brief, the channels were separated and the GFP-channel was used for quantification. Thus, background removal and smoothing were performed, a threshold was applied and the total area of the particles was determined.

### Cell viability analysis

The supernatant of treated cells was removed and cells were then lysed in 1x passive lysis buffer (5x passive lysis buffer diluted in deionized water; Promega, #E194A). For measuring metabolic activity, the CellTiter-Glo luminescent cell viability assay (Promega, #G7571) according to the manufacturer’s instructions was performed and the luminescent was measured by using an Orion II microplate Luminometer (Berthold) and the software Simplicity 4.2.

### Whole-cell and supernatant lysates

For preparing whole-cell lysates (WCL) collected cells were washed with DPBS, centrifuged for 4 min at 300 x g and the cell pellets were lysed in transmembrane lysis buffer supplemented with 1:500 protease inhibitor for 10 min on ice. Cell debris were pelleted by centrifugation for 20 min at 4 °C and 20,000 x g and the total protein concentration of the cleared lysate was determined using the Pierce Rapid Gold BCA Protein Assay Kit (Thermo Scientific, #A53225) according to the manufacturer’s protocol. The samples were adjusted to the same protein concentration with a transmembrane lysis buffer containing 1:500 protease inhibitor. For preparing cell-free lysates (CFL) cell debris of supernatants were removed by centrifugation for 4 min at 500 x g and the supernatants were transferred to fresh reaction tubes. After layering the supernatant on a cushion of 20% sucrose in PBS, samples were centrifuged for 90 min at 4 °C and 20,817 x g and the pellet was resuspended in transmembrane lysis buffer containing 1:500 protease inhibitor.

### SDS-PAGE and immunoblotting

SDS-PAGE and immunoblotting was performed as previously described^63^. In brief, whole-cell and supernatant lysates were mixed with 6x protein sample loading buffer (187.5 mM Tris-HCl prejusted to pH 6.8 (AppliChem GmbH, #A2264), 75% (v/v) glycerol (Sigma-Aldrich, #G5516), 6 % (w/v) SDS (Sigma-Aldrich, #8.22050), 0.3 % (w/v) Orange G (Sigma-Aldrich, #O3756), 15% (v/v) β-mercaptoethanol (Sigma-Aldrich, #444203) dissolved in deionized water) to a final concentration of 1x and heated up to 95 °C for 10 min before use. For protein separating, the samples were loaded on NuPAGE 4-12% Bis-Tris Gels (Invitrogen, #NP0321BOX) and the gels were running in 1x MES-SDS running buffer (20x MES-SDS running buffer diluted in deionized water; thermo scientific, #J62138.K2) for 90 min at 90 V. Next, the separated proteins were semi-dry blotted onto an Immobilon-FL PVDF-Membrane (Merck, #IPFL00010) at a constant voltage of 30 V for 30 min. After blocking in Blocker Casein in PBS (Thermo Scientific, #37528) for 1 h, the proteins on the membrane were stained with primary antibodies diluted in PBS-T (1x PBS with 0.2% (v/v) Tween 20 (Sigma-Aldrich, #P9416) and 0.1% (v/v) Blocker Casein in PBS) for 2 h at RT or overnight at 4 °C. Primary antibodies used in this study: Polyclonal rabbit anti-StrepII (1:2000; Abcam, #ab76949), monoclonal mouse anti-V5 (1:3000; Cell Signalling, #80076), monoclonal mouse anti-VSV-M (1:5000; Kerafast, #EB0011), polyclonal rabbit anti-LC3 (1:200; Sigma-Aldrich, #L8918), monoclonal mouse anti-p62 (1:1000; Abcam, ab56416), polyclonal rabbit anti-IFITM1 (1:500; Cell Signaling, #13126), polyclonal rabbit anti-IFITM2 (1:500; Abcam, #ab236735), monoclonal rabbit anti-IFITM3 (1:500; Cell Signaling, #59212), monoclonal rat anti-GAPDH (1:1000; Biolegend, #607902) and monoclonal mouse anti-β-actin (1:10,000; Sigma-Aldrich, #A5441). After incubation with the primary antibodies, the membrane was washed three times with PBS-T for 5 min at RT. Subsequently, the membrane was incubated in IRDye secondary antibodies diluted 1:20,000 in PBS-T. The following IRDye secondary antibodies were used in this study: IRDye 680RD Goat anti-Rabbit (LI-COR, #926-68071), IRDye 800CW Goat anti-Mouse (LI-COR, #926-32210), IRDye 680RD Goat anti-Rat (LI-COR, #926-68076) and IRDye 800CW Goat anti-Rat (LI-COR, #926-32219). After three times washing with PBS-T, the fluorescent signal of the secondary antibodies was detected using a LI-COR Odyssey (LI-COR) and the Image Studio Version 5.2 software. Image processing and quantification of band intensities were analyzed by using the software Image Studio Lite Version 5.0.21.

### siRNA-mediated knock down

1x 10^5^ HeLa-GL cells/well were grown in 500 µL medium in 24-well plates and one day later transfected with siRNA using Lipofectamine RNAiMax Transfection Reagent (Invitrogen, #13778150) and Opti-MEM reduced serum media according to the manufacturer’s instructions. For transfection the following siRNAs from Horizon Discovery ordered as SMARTPool were used: SNX12 (#M-013648-00-0005), STX12 (#M-018246-01-0005), TAB1 (#M-004770-02-0005), TMEM87b (#M-015008-00-0005) and ABCG2 (#M-009924-01-0005). As negative control Non-targeting Control siRNA#1 (Horizon Discovery, #D-001210-01-05) was used. Two days post transfection with siRNA, cells were either harvested for qRT-PCR analysis or transfected with plasmids encoding for E or E T9I harboring a StrepII tag (1 µg DNA/well) by using 2 µL 1x PEI and 100 µL Opti-MEM reduced serum media per well. In brief, 50 µL Opti-MEM were mixed with 2 µL 1x PEI and incubated for 5 min. Secondly, 50 µL Opti-MEM were mixed with 1 µg DNA. Afterwards both mixes were combined and after incubation (20 min, RT), the transfection mix was added to the cells. One day later the cells were further processed for Western Blot and autophagosome measurement by flow cytometry.

### Proximity Ligation Assay (PLA)

6x 10^4^ HeLa cells were seeded on glass cover slips in 24-well plates one day prior transient transfection using Lipofectamine3000 (Invitrogen, #L3000008) with plasmids coding for E or E T9I harboring a StrepII tag according to the manufacturer’s instructions. Cells were fixed 30 h post transfection with 3.7% PFA. Cell membranes were permeabilized with 0.5% Triton and blocked with 5% BSA (KPL, #5140-0006). PLA staining was performed as previously described^62,67^. Primary antibodies used: SNX12 Polyclonal Antibody (1:100; Invitrogen, #PA5-99046), STX12 Polyclonal antibody (1:100; Proteintech, #14259-1-AP), anti-TAB1 (1:100; abcam, #ab151408), TMEM87B Polyclonal antibody (1:100; Invitrogen, #PA5-57188), ABCG2 antibody (1:100; Santa Cruz Biotechnology, #sc-58222), rabbit anti-Strep-tag II antibody (1:450; abcam, #ab76949), mouse StrepII Tag Antibody (1:450; Novus Biologicals, #NBP2-43735). PLA probes and reagents used: Duolink In Situ PLA Probe Anti-Rabbit PLUS (Sigma-Aldrich, #DUO92002), Duolink In Situ PLA Probe Anti-Mouse MINUS (Sigma-Aldrich, #DUO92004), Duolink In Situ Detection Reagents FarRed (Sigma-Aldrich, #DUO92013).

### Molecular Modelling of SARS-CoV-2 E

The initial atomic positions were derived from the SARS-CoV-2 Envelope Protein Transmembrane Domain as reported in the 7k3g entry of the Protein Data Bank^68^. Equilibration at 300 K for 0.5 ns was performed by ReaxFF^68^ (reactive molecular dynamics) simulations using the Amsterdam Modeling Suite 2020 (http://www.scm.com). After equilibration, the amino acids of the Envelope Protein were replaced by the corresponding amino acids, together with the necessary modifications. Subsequently, an additional equilibration step (300 K for 0.5 ns) was performed by ReaxFF simulations in the NVT ensemble over 25 ps, with the system coupled to a Berendsen heat bath (held at T = 300 K with a coupling constant of 100 fs). Distances were calculated by averaging over these simulations. The program Visual Molecular Dynamics (VMD 1.9.3) was used for all visualizations^70^. The HOLE program was used to visualize the ion channel^71^.

### Phylogenetic and mutation abundance analyses

The phylogenetic trees were derived from Nextstrain^72^ at indicated timepoints. Abundance of the mutations in sequenced strains in the population was derived from data on Cov-spectrum^47^.

### Affinity purification and mass spectrometric analyses of A549 cells expressing E protein of SARS-CoV-2 strains

To determine the interactomes of E protein of SARS-CoV-2 strains, Strep-II tagged E proteins were used in four replicates each with eGFP as control. HEK293T cells were used to generate lentivirus for the Strep-II tagged E proteins. A549 cells (15 × 10^6^ cells per 15 cm dish) were transduced with lentiviral vectors encoding E proteins with 2.5 µg/ml puromycin selection. Cell pellets from two 15-cm dishes were used and lysed in lysis buffer (50 mMs Tris-HCl pH 7.5 (Trizma, Sigma Aldrich, #T1503), 100 mM NaCl (Sigma Aldrich, #S9888), 1.5 mM MgCl_2_ (Sigma Aldrich, #M8266), 0.2% (v/v) NP-40 (Sigma Aldrich, #I3021), 5% (v/v) glycerol (Sigma Aldrich, #49782), complete protease inhibitor cocktail (Roche), 0.5% (v/v) 750 U/μl Sm DNase). Further they were sonicated (15 min, 30 s on, 30 s off, high settings; Bioruptor, Diagenode). The protein lysates were normalized to 1 mg/ml for their concentration and subjected to affinity precipitation using 15 μl Strep-II tagged beads (IBA Lifesciences GmbH) with a constant agitation at 4 °C overnight. Subsequent washes with lysis buffer and washing buffer (50 mM Tris-HCl pH 7.5 (Trizma, Sigma Aldrich, #T1503), 100 mM NaCl (Sigma Aldrich, #S9888), 1.5 mM MgCl_2_(Sigma Aldrich, #M8266), 5% (v/v) glycerol (Sigma Aldrich, #49782)) were performed to remove non-specifically bound proteins. The enriched proteins were denatured, reduced, alkylated and digested by addition of 180 μl digestion buffer (0.6 M guanidinium chloride (Sigma Aldrich, #G3272)), 1 mM tris(2-carboxyethyl)phosphine (TCEP), 4 mM chloroacetamide (CAA) (Sigma Aldrich, #75259), 100 mM Tris-HCl pH 8, 0.5 μg LysC (WAKO Chemicals) and 0.5 μg trypsin (Promega) at 30 °C overnight at 300 rpm shaking. Three layers of C18 Empore filter discs (3M) were used to prepare the stage tips and peptide purification was performed. The purified peptides were subjected to LC-MS/MS. Peptides were loaded on a 20-cm reverse-phase analytical column (75 μm diameter; ReproSil-Pur C18-AQ 1.9 μm resin; Dr Maisch) and separated using an EASY-nLC 1200 system (Thermo Fisher Scientific). A binary buffer system consisting of buffer A (0.1% formic acid (FA) in H_2_O, Sigma Aldrich) and buffer B (80% acetonitrile (ACN#VWR), 0.1% FA (Sigma Aldrich) in H_2_O) with a 90-min gradient (5–30% buffer B (65 min), 30–95% buffer B (10 min), wash out at 95% buffer B (5 min), decreased to 5% buffer B (5 min), and 5% buffer B (5 min)) was used at a flow rate of 300 nl per min. Eluting peptides were directly analyzed on a Q-Exactive HF mass spectrometer in data-dependent acquisition (DDA) mode (Thermo Fisher Scientific)^15^. Data-dependent acquisition included repeating cycles of one MS1 full scan (300–1650 m/z, R = 60 000 at 200 m/z) at an ion target of 3 × 10^6^ with injection time of 20 ms. For MS2 scans the top 15 intense isolated and fragmented peptide precursors (R = 15 000 at 200 m/z, ion target value of 1 × 10^5^, and maximum injection time of 25 ms) were recorded. Dynamic exclusion, isolation window of the quadrupole, and HCD normalized collision energy were set to 20 s, 1.4 m/z, and 27 %, respectively.

### Data processing and analysis

Raw MS data files of AP–MS experiments conducted in DDA mode were processed with MaxQuant (version 1.6.14) using the standard settings and label-free quantification (LFQ) enabled (LFQ min ratio count 1, normalization type none, stabilize large LFQ ratios disabled). Spectra were searched against forward and reverse sequences of the reviewed human proteome including isoforms (UniprotKB, release 2019.10) and Strep-II tagged E proteins of SARS-CoV-2 strains and Strep-II tagged eGFP protein by the built-in Andromeda search engine^73^. Peptide and protein identification was controlled by a False Discovery Rate (FDR) of 0.01. Perseus was used to analyze the output of MaxQuant^74^. Protein groups identified as known contaminants or reverse sequence matches were excluded from the analysis. Only proteins with a minimum of two LFQ quantifications in at least one group of replicate experiments (*n* = 4) for a specific bait were considered for the analysis. Missing values were imputed using normal distribution, whose standard deviation was defined as 30% and the mean was offset by −1.8 s.d. of the data distribution of the real intensities observed in the corresponding mass-spectrometry run, respectively.

### Ethics

Approval for the Mount Sinai Pathogen Surveillance Program (MS-PSP) was obtained from the Mount Sinai Hospital (MSH) Institutional Review Board (IRB-13-00981)

### Statistical analyses

Statistical analyses were performed using GraphPad Prism 10. P-values were determined using a two-tailed Student’s t test with Welch’s correction. Unless otherwise stated, data are shown as the mean of at least three biological replicates ± SEM. Significant differences are indicated as: *, p < 0.05; **, p < 0.01; ***, p < 0.001. Unless otherwise specified, not significant (ns) differences are not indicated. Statistical parameters are further specified in the figure legends.

## DATA AVAILABILITY STATEMENT

The mass spectrometry proteomics data have been deposited to the ProteomeXchange Consortium via the PRIDE partner repository with the dataset identifier PXD048080. The data can be accessed with the following reviewer login information:

Username: Password: VMIgi7Zn

## ACKNOWLEDGEMENTS

We thank Daniela Krnavek, Martha Mayer, Kerstin Regensburger, Regina Burger, Jana-Romana Fischer and Birgit Ott for assistance. We thank Prof. Alexander Kühne (Ulm University) for providing access to the Leica SP5 microscope (DFG project ID: 432000323). We thank Dr. Antonio Piras (TUM University) for analyzing proteomics samples. We thank Klaus Klumpp, Gary Ewart and Michelle Miller (Biotron Limited) for providing the SARS-CoV-2 E inhibitor BIT225. The SARS-CoV-2 variants, B.1.617.2 (Delta) and B.1.1.529 (Omicron BA.5), were kindly provided by Prof. Dr. Florian Schmidt and Dr. Bianca Schulte (University of Bonn, Bonn, Germany). K.M.J.S. acknowledges support from the German Ministry for Research and Education (BMBF; IMMUNOMOD-01KI2014). This study was supported by DFG grants to K.M.J.S. (CRC1279-INST 40/623-1, SPP1923, SP 1600/6-1, SP 1600/7-1, SP1600/9-1), F.K. (CRC1279 and SPP1923) and A.P. (PI 1084/4, PI 1084/5 and TRR179/TP10, TRR237/A07, TRR353/B04). Work in the A.P.’s laboratory was supported by an ERC Consolidator grant (ERC-CoG ProDAP), the Helmholtz Association’s Initiative and Networking Fund (KA1-Co-02 “COVIPA”). D.K. is supported by a Baustein Grant from Ulm University, Else Kröner Fresenius Stiftung (2022_EKEA.47) and European Union’s Horizon 2020 Marie Sklodowska-Curie program (No. 101062524). S.K., H.H. and M.H. are part of the International Graduate School for Molecular Medicine, Ulm (IGradU). We further acknowledge the Bavarian State Ministry of Health, Bay-VOC (A.E.), IZKF Erlangen (Project A94) (A.E.) and the Bavarian Research Network FOR-COVID (A.P.).

