## Extended Data Fig. for "SARS-CoV-2 Omicron Envelope T9I adaptation confers resistance to autophagy"

**This PDF file includes:**

Extended data figures 1 to 5

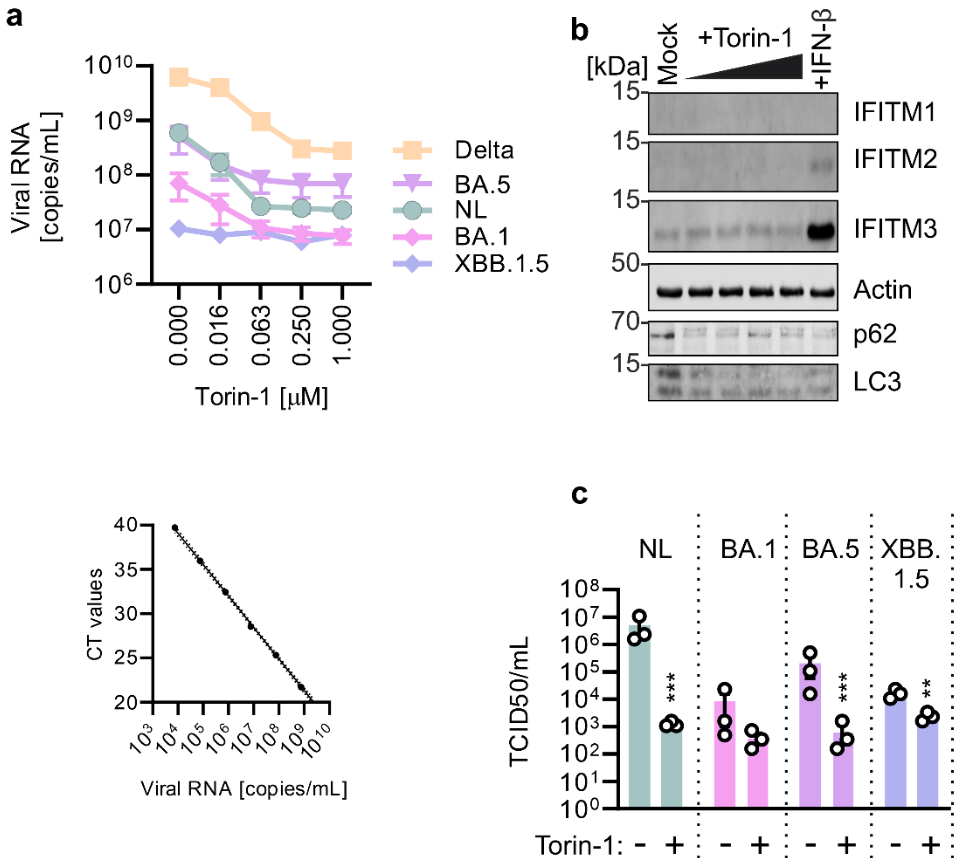


**Extended Data Figure 1.**

**Replication kinetics of NL-02-2020, Delta and Omicron upon autophagy induction and IFITM expression in Calu-3 cells. a,** Exemplary standard curve used for the quantification of viral RNA loads (left). Raw qRT-PCR values obtained from supernatants of Calu-3 cells infected with the indicated SARS-CoV-2 strains (MOI 0.05) for 48 h and treated with increasing amounts of Torin-1 (0.016-1 µM) (right). N = 3-6±SEM. **b,** Exemplary immunoblots of Calu-3 cells stimulated with increasing amounts of Torin-1 (0.016-1 µM), 1000 U IFN-β or medium for 24 h and stained with anti-IFITM1-3, anti-Actin, anti-p62, anti-LC3.**c,** Infectious SARS-CoV-2 in the supernatant of Calu-3 cells treated with 250 nM Torin-1 or left untreated and infected with indicated viruses (MOI 0.05) as assessed by TCID50. N = 3±SEM. Student's t-test with Welch's correction. **, $p<0.01$; ***, $p<0.001$.

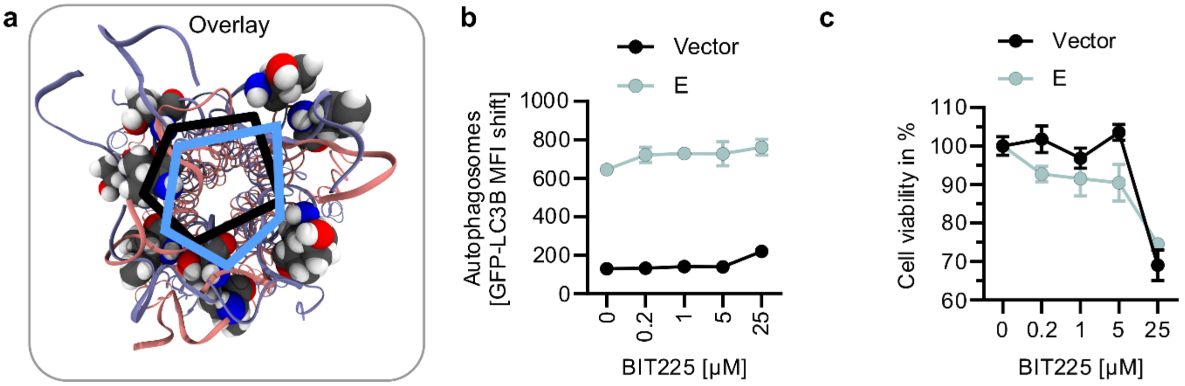


**Extended Data Figure 2.**

**Autophagy modulation by E is independent of E ion channel activity. a,** Visualization of the lateral projection surface of the E ion channel pore. Each case is an example image from the trajectory of the molecular dynamics simulation. **b,** Quantification of autophagosome levels by flow cytometry in HEK293T autophagy reporter cells (HEK293T-GL) transiently expressing SARS-CoV-2 E or a vector control and treated with increasing amounts of an E viroporin inhibitor BIT225 (0.2-25 µM) for 24 h. The measurement was assessed 48 h post transfection. N = 4±SEM. **c,** Effect of BIT225 treatment as in b on cell viability of HEK293T-GL, as assessed by intracellular ATP levels. N = 3±SEM.

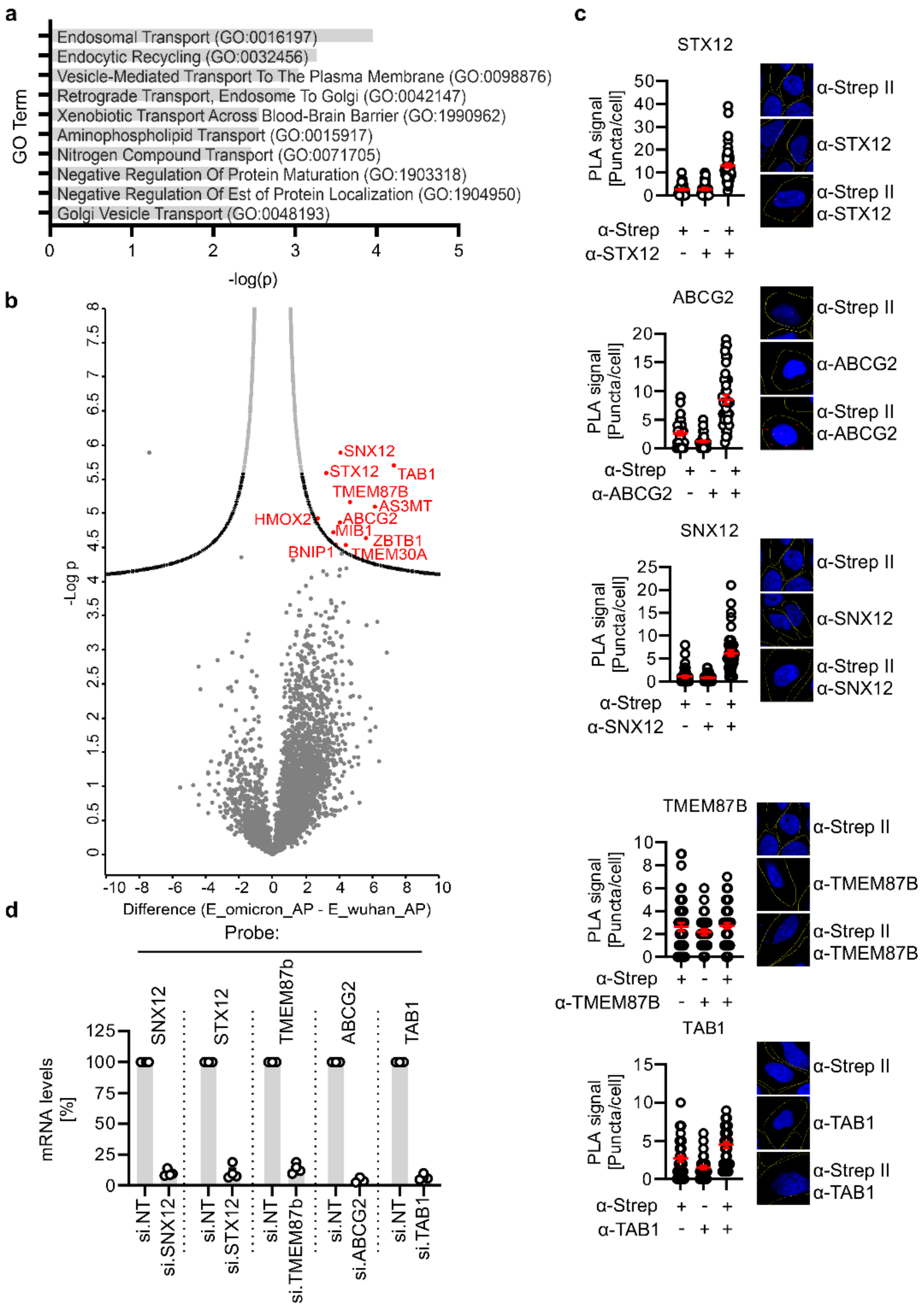


**Extended Data Figure 3.**

**E I9 has increased affinity to early autophagosome-associated proteins. a,** PantherDB aided GeneOntology analysis of the top100 hits from (Fig. 4b)**. b,** Volcano plot of the differential interactome analysis showing enriched proteins in E I9 pulldown versus the P value (-log P). Eleven highly significantly enriched proteins are highlighted in red and via labels**. c,** Single antibody controls of the Proximity ligation assays in Fig. 4c-g**. d,** Quantification of the mRNA levels of SNX12, STX12, TMEM87b, ABCG2 and TAB1 by qRT-PCR of HeLa-GL cells depleted of the indicated proteins by siRNA-mediated knock down (48 h post transfection). Data were normalized to the Non-targeting Control (100%). N = 4±SEM. Student's t-test with Welch's correction. *, p<0.05; **, p<0.01; ***, p<0.001.

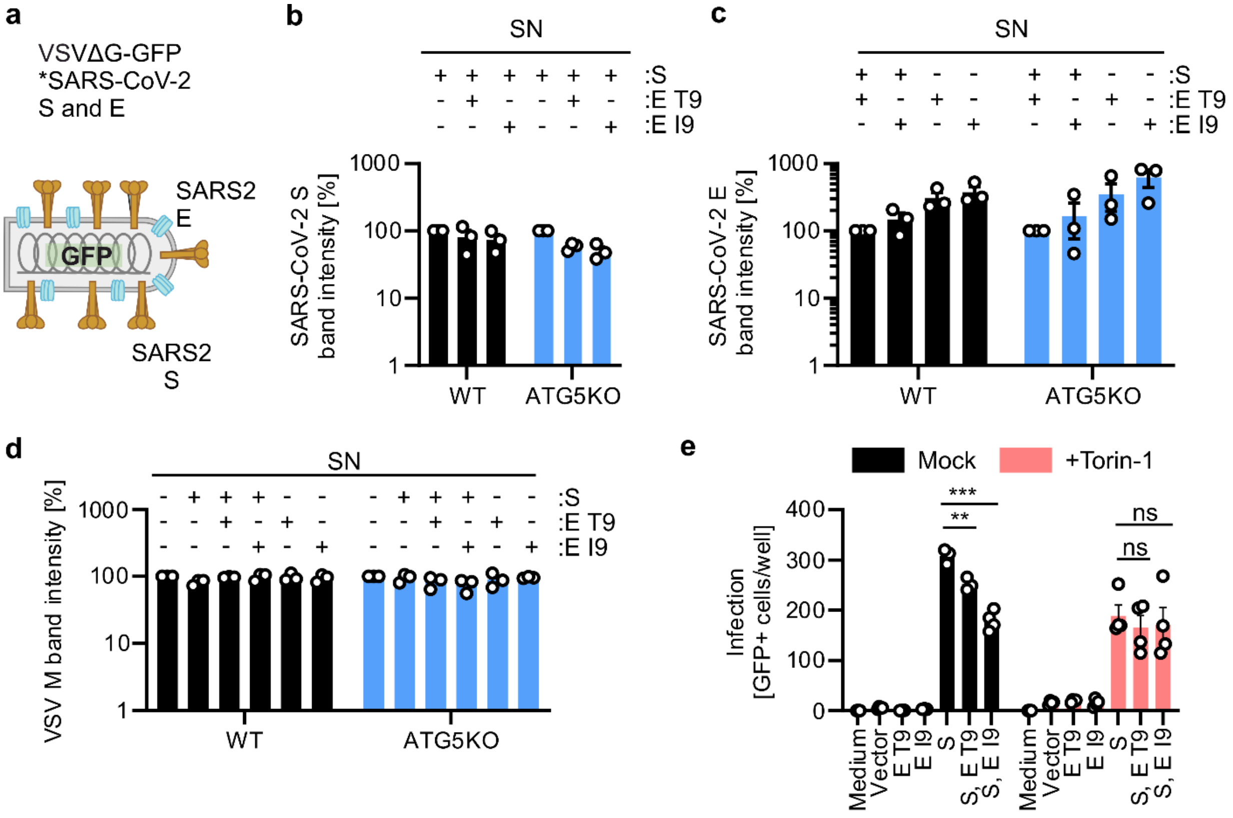


**Extended Data Figure 4.**

**Impact of E in pseudoparticle production and infection. a,** Schematic depiction of SARS-CoV-2 Spike (S) and Envelope (E) pseudotyped VSV-ΔG-GFP pseudoparticles. **b,** Quantification of the band intensities in (Fig. 5a) of S. S-only sample set to 100%. N = 3±SEM. **c,** Quantification of SARS-CoV-2 E band intensities from experiments in Fig. 5a. N = 3±SEM. **d,** Quantification of VSV-M band intensities from experiments in Fig. 5a. N = 3±SEM. **e**, Caco-2 cells treated with 0.5 μM Torin-1 or medium for 4 h before infection with VSV(GFP)ΔG pseudoparticles containing the indicated proteins. Infected GFP+ cells were automatically quantified after 22 h post infection. N = 4±SEM. Student's t-test with Welch's correction. *, $p<0.05$; **, $p<0.01$; ***, $p<0.001$. ns (non significant).

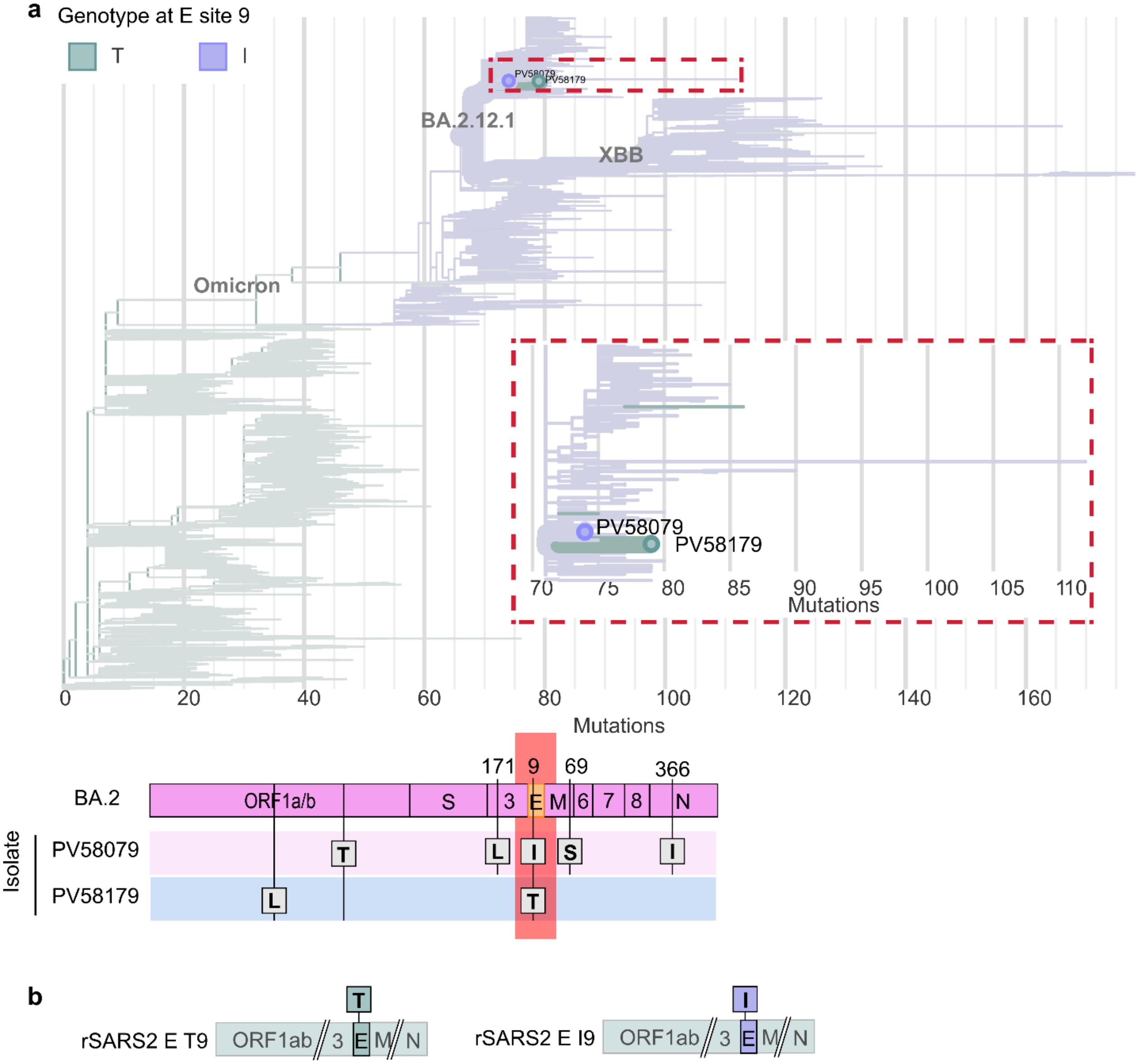


**Extended Data Figure 5.**

**Rare isolates and recombinant SARS-CoV-2. a,** Phylogenetic tree depicting the relation between the SARS-CoV-2 isolates PV58079 and PV58179. Mutations indicated are relative to HU-1. (bottom panel) Schematic depiction of patient isolate viruses PV58079 and PV58179 and their alterations compared to the BA.2 consensus sequence. **b,** Schematic depiction of recombinant SARS-CoV-2 (rSARS2) expressing either E T9 or E I9.
